# Improving the production and virulence of entomopathogenic fungi for biological control using insect-derived *in vitro* culture medium

**DOI:** 10.64898/2026.03.14.711814

**Authors:** AM Wilson, HH De Fine Licht

## Abstract

**Background:** The environment in which a fungus grows can directly influence their development, transmission, and pathogenic potential. This environment encompasses factors like nutrient availability, biotic and abiotic stressors, as well as host-derived chemical cues. In fungal pathogens, where conidia act as the infectious agents, the environment impacts the quantity and quality of these spores, thereby aOecting their ability to infect and kill hosts. In the present study, we investigated the effect of host-derived medium types on various phenotypes, including spore production, growth rate, and virulence in two entomopathogenic fungi, *Metarhizium acridum* and *Metarhizium brunneum*. Three medium types derived from insect material were compared to a standard laboratory medium.

**Results:** Conidia produced on the insect-derived media exhibited enhanced sporulation and reduced time to sporulation, while conidial germination and maximum growth rate were comparable across medium types, suggesting that some of the medium-induced phenotypic effects were transient. Notably, conidia derived from two of the insect medium types demonstrated higher virulence, indicating that host-derived cues may prime virulence.

**Conclusion:** These results highlight that the composition of growth substrates can regulate fungal reproductive strategies and virulence, with implications for developing high-throughput phenotyping and for the biotechnological optimization of mass production and efficacy of entomopathogenic fungi in biological control applications.

**GRAPHICAL ABSTRACT:** 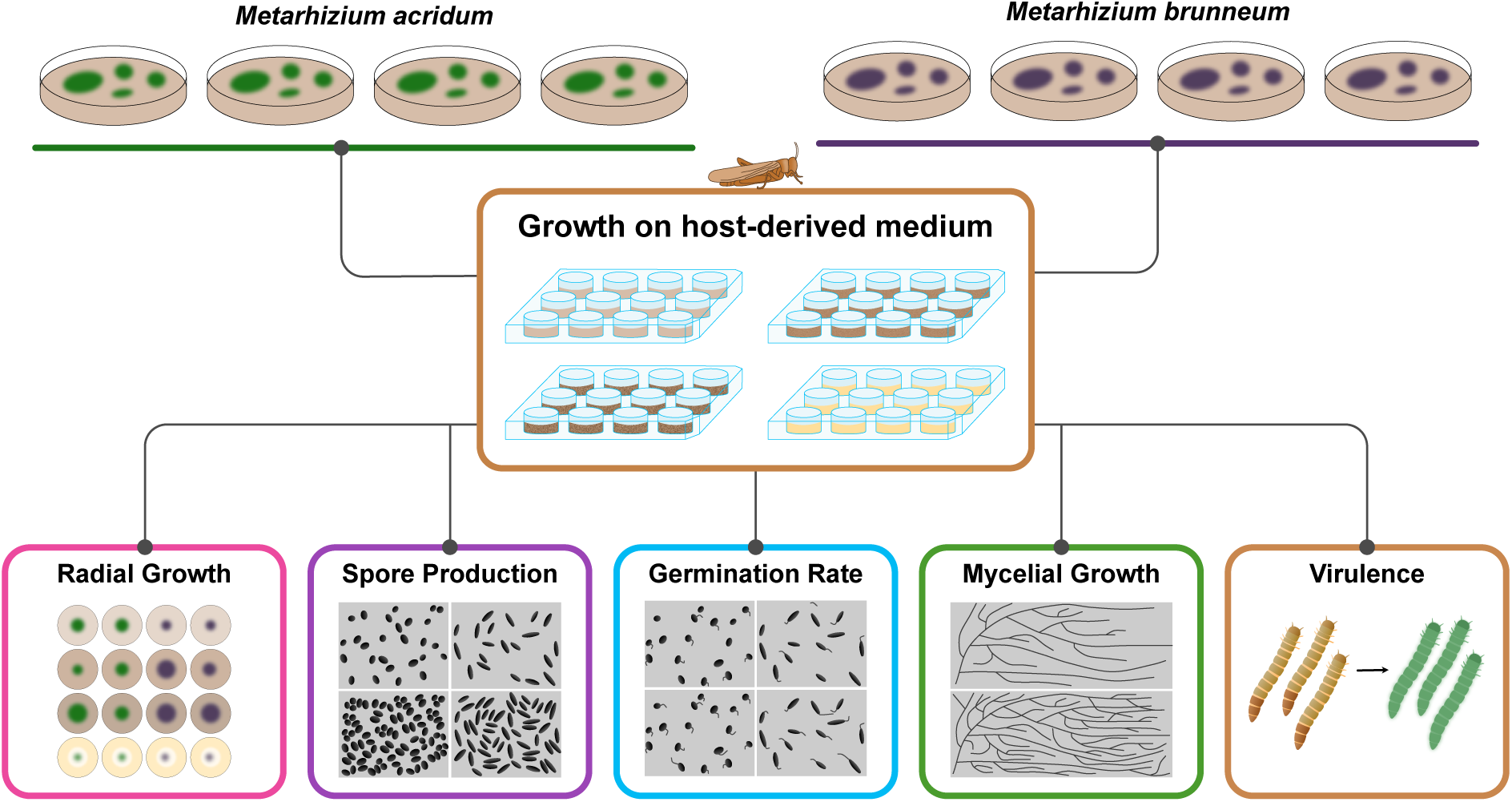

## BACKGROUND

Entomopathogenic fungi play an important role in the suppression of insects in natural ecosystems (Augustyniuk-Kram and Kram, 2012). This ecological function can be exploited in agricultural settings and in human health, where species like *Metarhizium anisopliae* and *Beauveria bassiana* can be employed as biopesticides, combating agricultural pests like thrips and aphids (Smagghe et al., 2023) and human disease vectors like mosquitoes (Lovett et al., 2019; Shoukat et al., 2020). Given the increasing demand for environmentally sustainable alternatives to chemical pesticides, entomopathogenic fungi have been the focus of extensive research into improving their mass production and performance (Iwanicki et al., 2023, 2021; Jaronski and Mascarin, 2017; Mascarin et al., 2024).

For most entomopathogenic fungi, conidia- the asexual spores- act as the infectious agent (Quesada-Moraga et al., 2022; St. Leger, 2024). It is this life stage that adheres to the host’s cuticle, germinates, and results in host penetration and subsequent colonization. Once the host’s internal resources have been fully consumed, the fungus will produce dispersible and infectious conidia on the external surface of the host, allowing the lifecycle to restart on a new host (De Fine Licht et al., 2025). The quality, viability, longevity, and yield of these spores are thus important factors to consider when optimizing the production of entomopathogens for biocontrol systems (Jaronski and Mascarin, 2017). Importantly, these traits are not fixed and can be significantly impacted by the environment in which the spore is produced (Sala et al., 2023). Thus, there is often a trade-off between rapid fungal growth in industrial production environments and the efficacy of infectious conidia in the field.

Species in the fungal genus *Metarhizium* are known to be influenced by the culture in which they are grown (De Fine Licht et al., 2025). The impact of the growth medium’s nutrient composition has been studied in species such as *M. anisopliae* (Ibrahim et al., 2002; Shah et al., 2005)*, M. robertsii* (Iwanicki et al., 2018), and *M. flavoviride* (Fargues et al., 2001) where it has been shown that the medium can affect phenotypes like spore production, germination, adhesion, appressorial development, and virulence. It is thus clear that culture medium impacts the production of spores as the fungus is actively using the available nutrients, but growth conditions can also impact the quality and viability of the spores in the longer term. For example, the C:N ratio of the conidia is partly determined by the medium of origin and this ratio can have long last effects on the viability, stress tolerance, and virulence of the conidia (Shah et al., 2005). However, most investigations into the impact of growth medium of entomopathogenic fungi have focused on the standard artificial media most commonly used for the maintenance and propagation of these species (Fargues et al., 2001; Shah et al., 2005), the use of agro-industrial waste products (Sala et al., 2023, 2021), and the need for a solid fermentation medium for conidia production (Loera et al., 2016). Knowledge regarding the effect of host-derived nutrients and cues on the production and viability of conidia is thus lacking.

Entomopathogenic fungi like members of the genus *Metarhizium* alter their physiology and morphology in response to specific insect hosts (Iwanicki et al., 2022, 2020, 2025). The specific attributes of the variable tissues and organs of the insect host body also influence the physiology of invading entomopathogenic fungi (De Fine Licht et al., 2025; Shik et al., 2025). It is thus clear that the quantity and composition of specific insect-derived nutrients and other biotic cues could impact entomopathogenic fungi. However, little is known regarding the impact of host derived nutrients and cues on the biology, and ultimately conidia production, of these species. Notably, being able to utilize any positive effects of insect-derived nutrients and chemical cues for high-throughput phenotypic screening or large-scale production of entomopathogenic fungi for biological control purposes could potentially improve the efficacy of solid-state fermentation production of conidia.

The primary aim of this study was to determine whether insect-derived medium types influence phenotypes such as sporulation, growth, germination, and virulence in the entomopathogenic fungi *Metarhizium acridum* and *Metarhizium brunneum*. This was achieved by comparing these phenotypes of fungal isolates grown on a standard growth medium to isolates grown in an artificial medium supplemented with different locust-derived materials, an insect host species suitable for both the locust-specialist, *M. acridum* and the insect-generalist, *M. brunneum*.

## METHODS

### 1. Media types used

Four different types of solid media (A – D) were used (**Figure 1**, **Table 1**). Three of these were produced by supplementing minimal media (**Table 2**) with 20g/l of freeze-dried and ground adult *Locusta migratoria* (∼6-8 cm length). The adult locusts were freeze-dried in a ScanVac CoolSafe freeze drier (LaboGene, Allerød, Denmark) at −55°C for up to 48 hours, after which they were ground to a powder using a MultiDrive basic industrial blender (IKA, Staufen, Germany). Care was taken to ensure that all remaining large and intact pieces were removed, resulting in a mixture consisting of a particle size smaller than 5mm in length and width.

**Figure 1:**
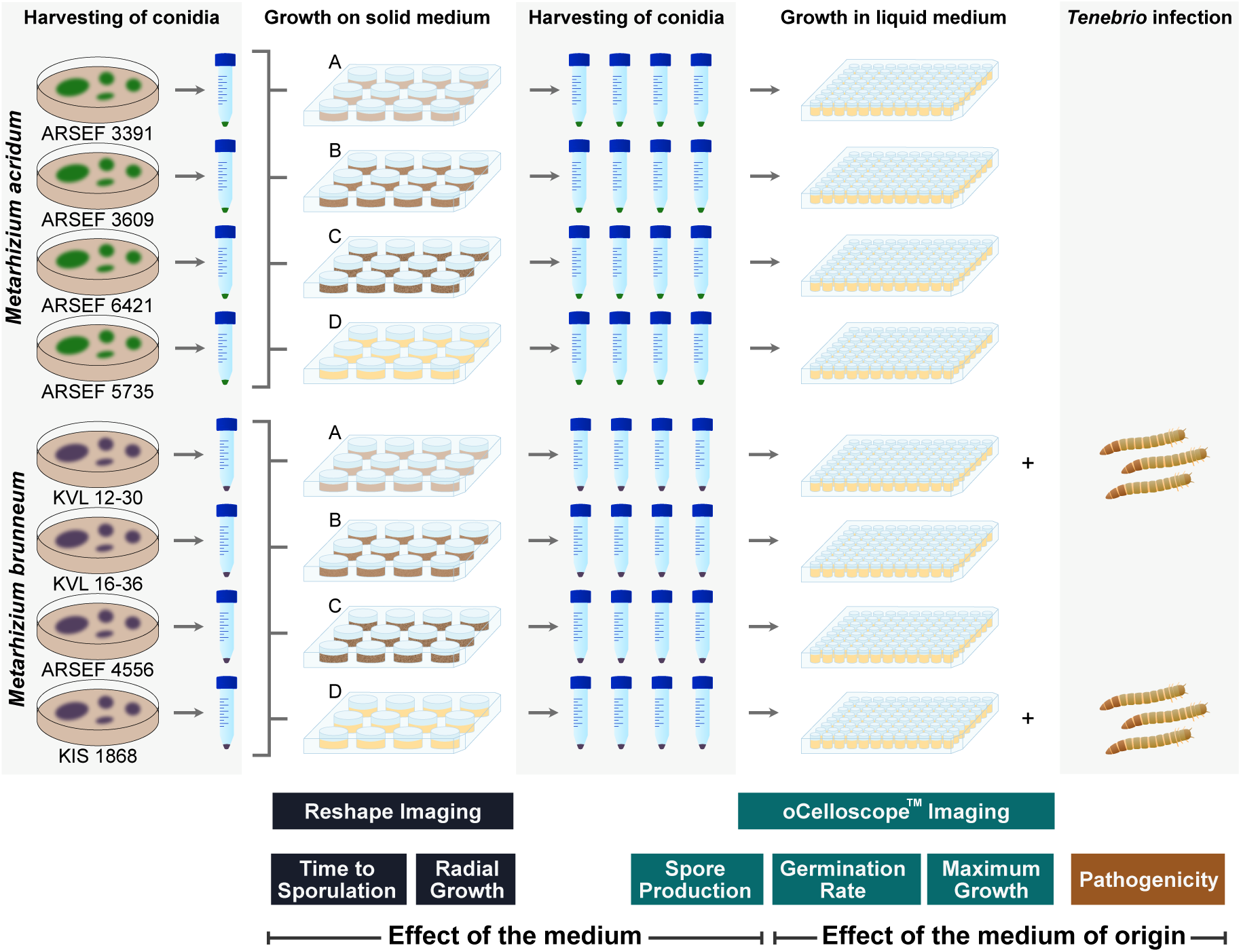
The experimental setup used to assess the short- and long-term effects of growing entomopathogenic fungi on media derived from host tissue. Four isolates of each species were used and grown on three locust-derived media (A – C) and the ¼ SDAY control medium (D). Medium A represents the twice filtered locust medium, Medium B represents the once filtered locust medium, and Medium C represents the unfiltered locust medium. Over the course of 7 days, radial growth and sporulation were monitored using a Reshape imaging robot. Thereafter, conidia were harvested to measure spore count, germination rate, and maximum growth in ¼ SDY medium using an oCelloscope^TM^ imaging system. Lastly, pathogenicity assays were conducted using selected *M. brunneum* isolates on using *Tenebrio molitor* larvae.

**Table 1.**
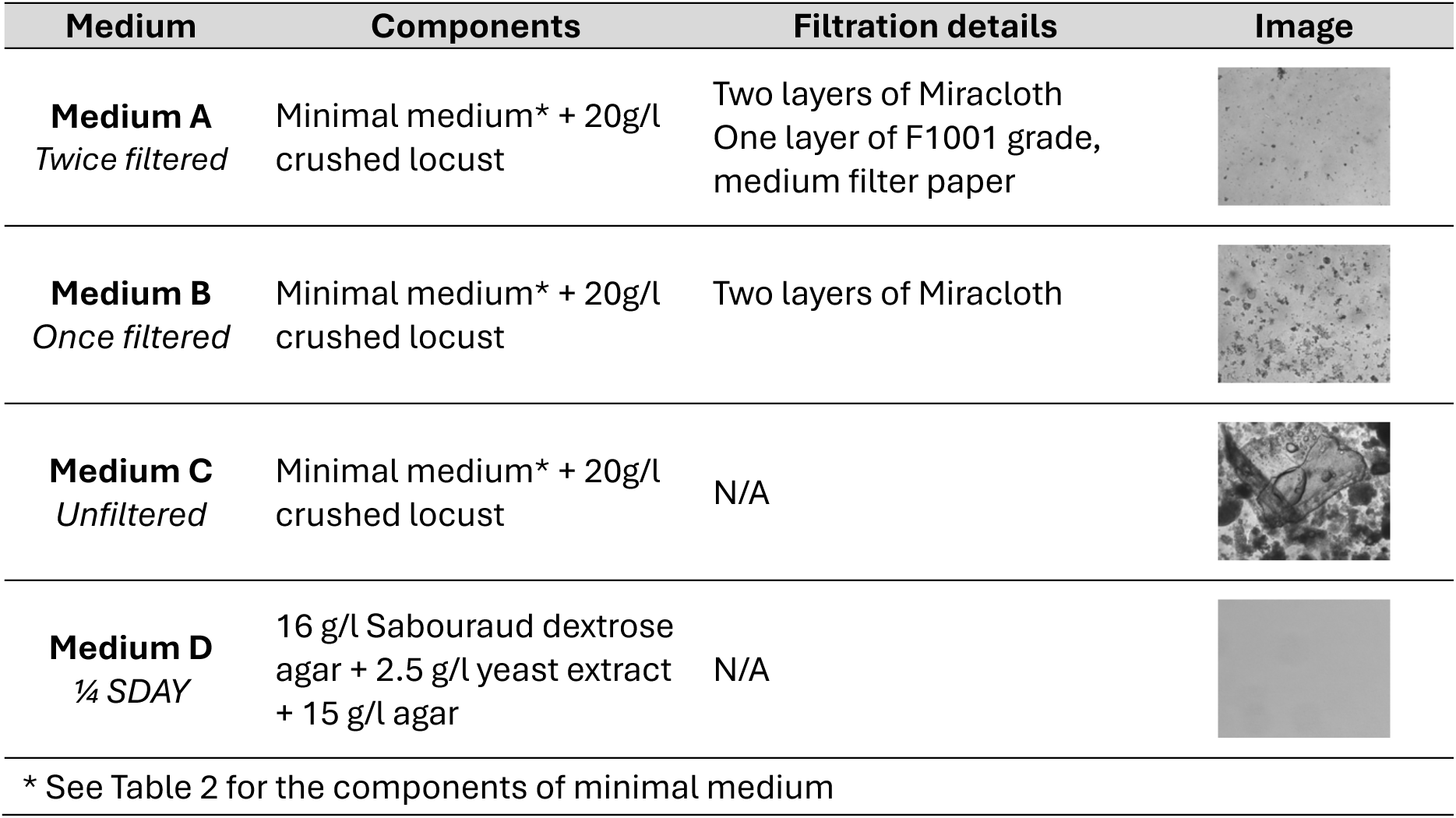
Details regarding the four media types used in this study.

| Medium | Components | Filtration details | Image |
| --- | --- | --- | --- |
| **Medium A**<br>*Twice filtered* | Minimal medium* + 20g/l crushed locust | Two layers of Miracloth<br>One layer of F1001 grade, medium filter paper |  |
| **Medium B**<br>*Once filtered* | Minimal medium* + 20g/l crushed locust | Two layers of Miracloth |  |
| **Medium C**<br>*Unfiltered* | Minimal medium* + 20g/l crushed locust | N/A |  |
| **Medium D**<br>*¼ SDAY* | 16 g/l Sabouraud dextrose agar + 2.5 g/l yeast extract + 15 g/l agar | N/A |  |

\* See Table 2 for the components of minimal medium

**Table 2.** Components of the minimal medium used as a base for the three locust media types.

| Component | Chemical formula | Quantity |
| --- | --- | --- |
| Monopotassium phosphate | $\text{KH}_2\text{PO}_4$ | 0.4 g |
| Disodium hydrogen phosphate dihydrate | $\text{Na}_2\text{HPO}_4 \cdot 2\text{H}_2\text{O}$ | 2 g |
| Magnesium sulfate heptahydrate | $\text{MgSO}_4 \cdot 7\text{H}_2\text{O}$ | 0.6 g |
| Potassium chloride | KCl | 1 g |
| Potassium nitrate | $\text{KNO}_3$ | 0.7 g |
| Agar |  | 15 g |
| Water |  | Up to 1000 ml |

Medium A (also referred to as “twice filtered locust medium”) was a locust medium that had been filtered through two layers of Miracloth (Millipore Sigma) and one layer of F1001 grade, medium filter paper (CHMLAB Group, Spain) to produce an almost-transparent locust-derived medium with few particulates. Medium B (also referred to as “once filtered locust medium”) was a locust medium that was only filtered through two layers of Miracloth. Medium C (also referred to as “unfiltered locust medium”) was not filtered and retained all of the larger locust particulates. All three of these media were prepared by combining the minimal medium reagents (**Table 2**) and crushed locust before autoclaving at 121°C for 20 min. Once cooled to room temperature, Medium A and B were filtered as described above. Agar was then added to all three media, which were then autoclaved as before. Medium D (also referred to as “¼ SDAY”) was prepared as the control medium (**Table 1**), as this is a medium frequently used for the growth and maintenance of entomopathogenic fungi. Media A – D were poured in 24-well culture plates, with approximately 2 ml per well and stored at room temperature before use. Lastly, ¼ SDY broth was also prepared.

### 2. Isolates used

Two species of entomopathogenic fungi were used for this study: *Metarhizium acridum*, a locust specialist, and *Metarhizium brunneum*, an insect generalist. Four isolates of both species were used (Moustaka et al., 2021; Parker et al., 2023; Razinger et al., 2018; Steinwender et al., 2014) (**Table 3**). These isolates were retrieved from storage as glycerol spore suspensions at −70°C and plated onto unfiltered locust medium. These fresh cultures were stored in the dark at 25°C for up to 10 days, after which spores were harvested using a 0.05% Triton X-100 solution and standard procedure. Spore suspensions of 1×10^6^ spores/ml were prepared and used for the growth and sporulation assays.

**Table 3.** Isolates used in this study.

| Species | Isolate | Origin, Host, Date |
| --- | --- | --- |
| *M. acridum* | ARSEF 3391 | Tanzania, *Zonocerus elegans*, Rec. 1991 |
| *M. acridum* | ARSEF 3609 | Thailand, *Patanga succincta*, Rec. 1992 |
| *M. acridum* | ARSEF 6421 | Senegal, *Kraussaria angulifera*, Rec. 1999 |
| *M. acridum* | ARSEF 5735 | Madagascar, Orthoptera, Rec. 1998 |
| *M. brunneum* | KVL 12-30 | Denmark, soil (baited using *T. molitor*), 2014 |
| *M. brunneum* | KVL 16-36 | Pure cultures obtained from Met52 Granular (Novozymes) |
| *M. brunneum* | ARSEF 4556 | United States of America, *Boophilus* sp. 1993 |
| *M. brunneum* | KIS 1868 | Slovenia, *Agriotes* sp. 2018 |

### 3. Growth and sporulation assays

To determine the effect of medium type on fungal growth rate (radial growth) and spore production (number of conidia produced), these phenotypes were assayed on solid media A - D. A total of 5 μl of each spore suspension was inoculated onto the four media types, in triplicate. The 24-well plates were transferred into Reshape Robot imaging system (Reshape Biotech, Denmark) and incubated at 25°C for 7 days. Images were captured every 2 hours.

The images generated by the Reshape Robot imaging system were used to assess average growth rate and estimated time to sporulation. Average radial growth was estimated at hour 60 (image 31) and hour 120 (image 61) by measuring the diameter of each culture on two perpendicular axes using ImageJ v1.54g. These values were used to calculate a per day growth rate. Where possible, time to sporulation was assessed by visually inspecting each image and identifying evidence of sporulation. This included the darkening or greening of otherwise white mycelium which later led to the confirmed production of conidia.

Spore production was measured by calculating the total number of spores produced by each isolate on each medium type after a 7-day incubation. The spores were harvested by adding 2ml of a 0.5% Triton X-100 solution to each culture and firmly scraping with an inoculation loop to release the spores. A total of 1ml of the resulting spore suspension was filtered through a 70μm pore cell strainer (VWR, Avantor Science Central, Denmark) and used to produce serial dilutions. Spore counting was performed on each of the dilutions using an oCelloScope Imaging system (BioSense Solutions, Denmark) and the UniExplorer v 14.3 software, with the *Metarhizium* algorithm for spore counting and a sensitive threshold of 0.3.

### 4. Germination and growth assays

To determine whether longer lasting phenotypic effects were carried over in spores that were produced on different medium types, the germination and subsequent mycelial growth of these spores were assayed in liquid medium. Each of the replicates from above were pooled and spore suspensions of 5 × 10^5^ spores/ml were prepared per isolate. These suspensions were combined with ¼ SDY medium to a final concentration of 5 × 10^4^ spores/ml and 100 μl was transferred into a 96-well plate. Each isolate x medium of origin combination was plated in triplicate and the oCelloScope Imaging system (BioSense Solutions, Denmark) was used to track the germination and growth of the isolates over the course of 48 hours. Images were captured every hour, with an image distance of 4.9 μm and a scan area length of 405 μm. The UniExplorer v14.3 (BioSense Solutions, Denmark) spore counting algorithm was used to estimate germination rate at 12 and 24 hours. The same software was used to calculate the growth rate as fungal pixels in each image per time unit, using the filamentous growth algorithm, which identifies the five consecutive data points with the steepest slope on the linear growth curve of log-transformed pixel-data to estimate the maximum growth rate similar to (Hall et al., 2014).

### 5. Virulence bioassay

Virulence assays were performed using conidia from two of the *M. brunneum* isolates, KVL 12-30 and KVL 19-39, on *Tenebrio molitor* larvae (Coleoptera, Tenebrionidae). These two isolates were chosen as they produced a sufficient quantity of conidia on all four medium types. Isolates of *M. acridum* were not used for this assay as they do not naturally infect *T. molitor*. At the time of infection, the larvae weighed an average of 53 ± 4.1 mg and were being maintained at 23 °C in the dark on wheat bran with slices of organic potato. The virulence bioassay was conducted in three repetitions with 10 larvae per treatment (isolate x medium combination).

Spore suspensions of 10^7^ spores/ml were prepared per treatment. The larvae were deposited into the suspensions and gently swirled for 10 s before being transferred to sterile filter paper for 5 mins. Control samples were treated with a 0.005% Triton X solution. The 10 larvae per treatment were transferred into a 180 ml plastic cup with a ventilated lid and supplied with a 2 g slice of organic potato to increase humidity and allow for fungal germination and penetration. After 24 hrs, the potato slice was replaced with 1.5 g of wheat bran and a fresh potato slice. The wheat bran and potato were replenished as needed every 2-3 days. The larvae were maintained at 23 °C in the dark for 10 days and were inspected every 24 hrs for mortality. The dead larvae were removed, surface sterilized in 5% sodium hypochlorite, and washed twice in sterile water. The cleaned cadavers were placed in 30 ml plastic cups with moist filter paper and monitored for fungal development.

### 6. Statistical analyses

Statistical analyses were performed using R 4.5.1 (The R Foundation for Statistical Computing 2025) and the `tidyverse` package (Wickham et al., 2019), and plots were generated using the `ggplot2` package (Wickham, 2016). For spore production, time to sporulation, radial growth, conidial germination, and maximum linear growth rate, univariate analyses were performed on all measured traits. Either linear models or generalized linear models were used depending on the distribution of each trait (**Table S1**). The medium type, isolate, and their interaction were included as fixed effects in all models.

The significance of each of these factors on the measured phenotypes was assessed using Type II ANOVA tests implemented in the `Anova` function of the `car` package (Fox and Weisberg, 2018), after checking for normality, independence, and homoscedasticity where relevant. F-tests were used for the Gaussian linear models and likelihood-ratio χ^2^ tests were used for generalized linear models. When a significant result was identified, the `emmeans` package (Russel, 2023) was used to estimate marginal means and compute pairwise comparisons, including a false discovery rate (FDR) correction for multiple testing. Pairwise comparisons were displayed using the `cld` function in the `multcomp` package (Hothorn et al., 2008).

Larval survival was analysed using time-to-event methods, with survival time being defined as the number of days from the start of the experiment until the day that death was recorded. Larvae that were alive on day 10 were treated as right-censored observations. Survival curves were estimated using the Kaplan-Meier method, with differences in survival among treatments being assessed through a log-rank test. To quantify differences in mortality risk, Cox proportional hazard (coxph) regression models were fitted with medium, isolate, medium/isolate interaction, and repetition as fixed effects: Medium + Repetition (**Table S1**). Hazard ratios (HR) with 95% confidence intervals (CI) were calculated. Control samples were included in Kaplan–Meier survival plots to confirm absence of background mortality but were excluded from coxph models due to zero mortality. These analyses were performed using the `survival` and `survminer` R packages (Kassambara et al., 2026; Lin and Zelterman, 2002).

## RESULTS

### 1. Medium influences spore production in both entomopathogenic fungi

The average spore count was significantly affected by medium (χ^2^ = 145.42, df = 3, p < 0.001), isolate (χ^2^ = 142.78, df = 3, p < 0.001), and the interaction between the two (χ^2^ = 197.64, df = 9, p < 0.001) in *M. acridum* (**Figure 2**, **Table S1 & S2**). For three of the isolates, more spores were produced on the unfiltered locust medium than the ¼ SDAY control medium, suggesting that the presence of host-derived nutrients and other cues increases spore production. The effect of filtering the locust medium had varied effects on spore production, depending on the isolate. For example, in ARSEF3391, the twice filtered medium resulted in spore production lower than the other two locust media types, while in ARSEF3609, filtration had no effect on spore production.

**Figure 2:**
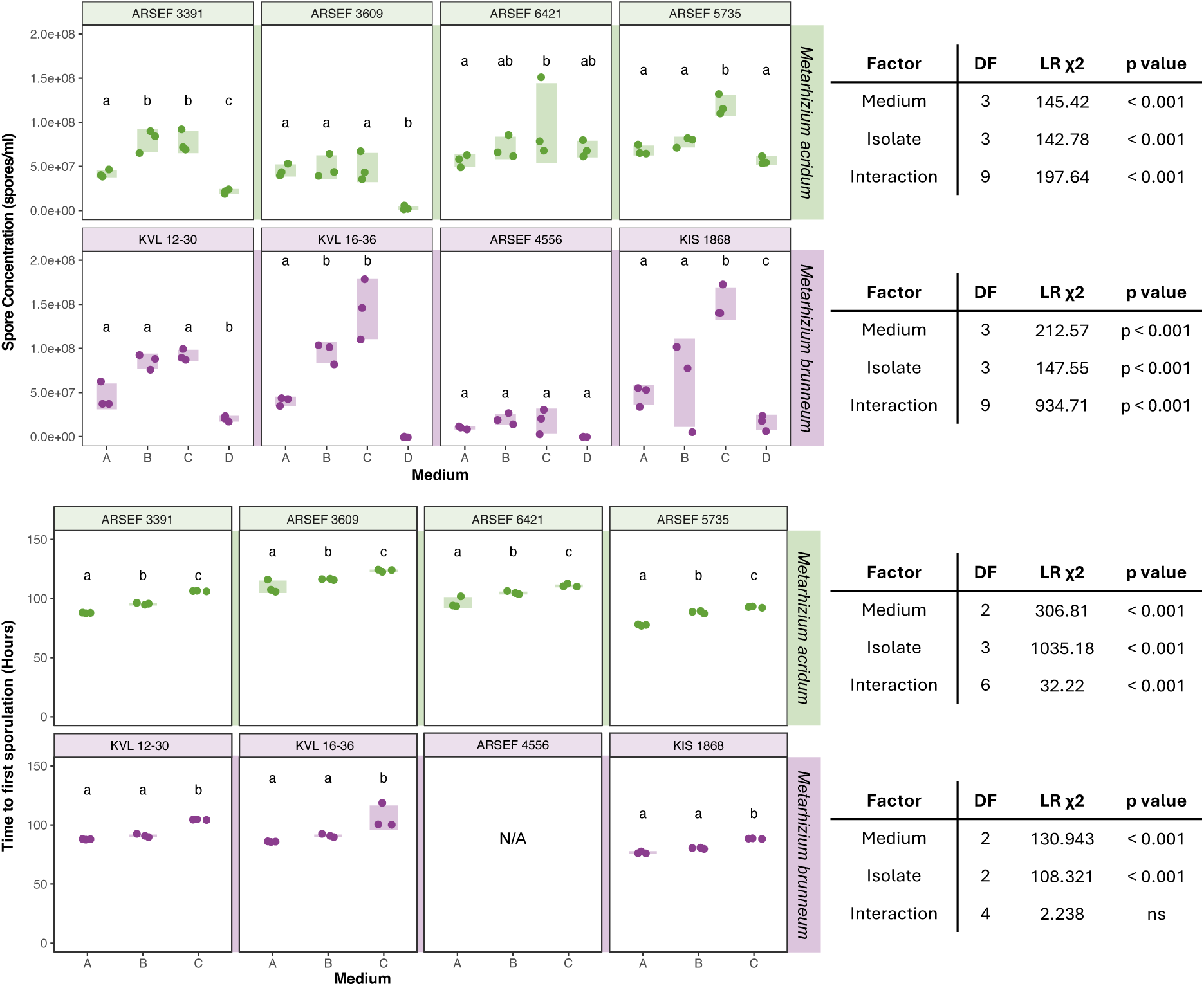
The effect of medium type on spore production in *Metarhizium acridum* and *Metarhizium brunneum*. In both species, at least one of the locust-derived medium types results in the production of a higher number of spores than the ¼ SDAY medium, a culture medium that is commonly used for the maintenance of entomopathogenic fungi. This was particularly pronounced in *M. brunneum*. Time to first sporulation was also significantly impacted by the medium type, with the early stages of sporulation typically not being visible in the 1/4 SDAY medium but visible as early as before 80 hours post-inoculation. In all plots, the shades area behind the plotted points represents the data’s spread from minimum to maximum values.

Spore production was similarly affected in *M. brunneum* (**Figure 2**, **Table S1 & S2**), with medium (χ^2^ = 212.57, df = 3, p < 0.001), isolate (χ^2^ = 147.55, df = 3, p < 0.001), and the interaction between the two (χ^2^ = 934.71, df = 9, p < 0.001) significantly influencing spore production. Filtration had a variable effect on the spore production. For example, in KVL1230, spore production wasn’t affected by filtration, but in KVL1636, the cultures grown on twice filtered medium produced significantly fewer spores than the cultures grown on the two other locust media. Notably, no spores could be detected in KVL1636 and ARSEF4556 in the ¼ SDAY control medium after 7 days of incubation.

The cultures were visually inspected to identify when the first evidence of sporulation was detectable. In most cases, it was not possible to confidently identify spore production on the ¼ SDAY control medium within the seven days of monitoring, but it was possible on all three locust media. The time to first sporulation was significantly influenced by the medium (χ^2^ = 306.81, df = 2, p < 0.001), isolate (χ^2^ = 1035.18, df = 3, p < 0.001), and the interaction between the two (χ^2^ = 32.22, df = 6, p < 0.001) in *M. acridum*, with the twice filtered locust medium resulting in the shortest time to sporulation (**Figure 2**, **Table S1 & S2**). In *M. brunneum*, the time to first sporulation was significantly impacted by the medium (χ^2^ = 130.943, df = 2, p < 0.001) and the isolate (χ^2^ = 108.321, df = 2, p < 0.001). In this case, both the once and twice filtered locust media resulted in the earliest onset of sporulation (**Figure 2**, **Table S1 & S2**). The *M. brunneum* isolate ARSEF4556 was excluded from this analysis as evidence for sporulation was only detectable on the unfiltered locust medium.

### 2. Radial growth is diZerently aZected by medium in the two entomopathogens

The average daily growth rate was significantly affected by medium (F_3,32_ = 40.00, p < 0.001), isolate (F_3,32_ = 14.93, p < 0.001), and the interaction between the two (F_3,32_ = 4.01, p < 0.01) in *M. acridum* (**Figure 3**, **Table S1 & S2**). In general, the four isolates grew fastest on the unfiltered locust medium and slowest on the twice filtered locust medium. In contrast, the medium affected the four *M. brunneum* isolates in vastly different ways (**Figure 3**, **Table S1 & S2**). While medium (F_3,32_ = 31.22, p < 0.001), isolate (F_3,32_ = 31.70, p < 0.001), and the interaction between the two (F_3,32_ = 12.17, p < 0.001) impacted growth, pairwise differences were only seen in KVL1636 and ARSEF4556, where the ¼ SDAY control medium elicited the quickest growth. In contrast, there were no significant differences in growth when KVL1230 and KVL1939 were grown on the different medium types.

**Figure 3:**
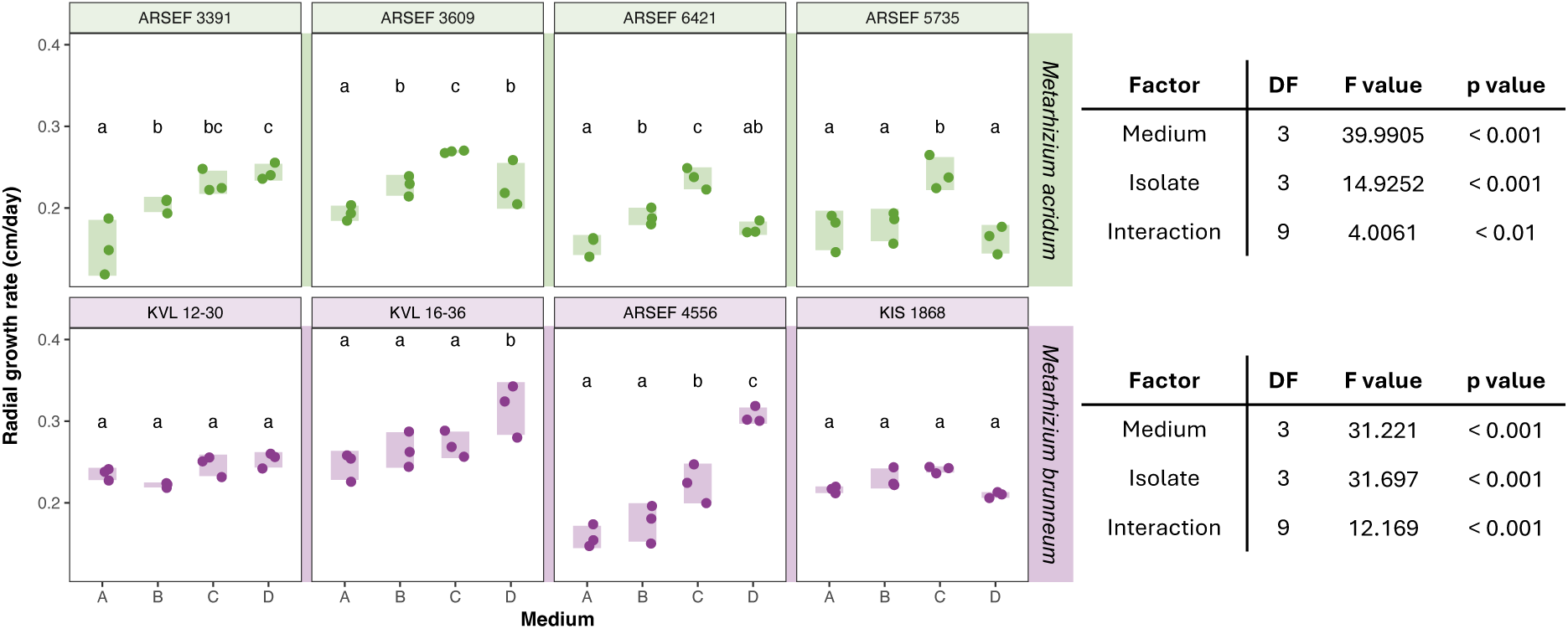
The effect of medium type on radial growth in *Metarhizium acridum* and *Metarhizium brunneum*. Radial growth in *M. acridum* was typically quickest on the unfiltered locust medium and slowest on the twice filtered locust medium, suggesting that *M. acridum* benefits from the additional nutrients present in unfiltered host-derived media. In contrast, growth was only impacted by the medium in two of the *M. brunneum* isolates, with minor differences seen between the three locust-derived media types, and the ¼ SDAY medium resulting in the quickest growth in these two isolates. In all plots, the shades area behind the plotted points represents the data’s spread from minimum to maximum values.

### 3. Medium of origin impacts the two species diZerently

In general, the medium of origin did not have an effect on germination in ¼ SDAY medium at 12 hours (**Figure 4**, **Table S1 & S2**). The exception was *M. acridum* isolate ARSEF3609, which showed a slightly higher germination rate at 12 hours when the conidia originated from unfiltered locust medium. The medium of origin also had no impact on the maximum growth rate of the mycelium in liquid medium for *M. acridum* (F_3,32_ = 1.64, ns). In *M. brunneum*, the medium (χ^2^ = 73.59, df = 3, p < 0.001), the isolate (χ^2^ = 27.08, df =1, p < 0.001), and their interaction (χ^2^ = 22.41, df = 3, p < 0.001) significantly influenced germination rate. The conidia produced on the ¼ SDAY control medium showed lower rates of germination than those produced on the three locust media for both isolates analysed. In contrast, the medium of origin had no impact on the maximum growth rate of the mycelium in liquid medium for *M. brunneum* (F_3,16_ = 0.25, ns). Because KVL1636 and ARSEF4556 did not produce countable spores on the ¼ SDAY control medium, germination and growth could not be assessed for these isolates.

**Figure 4:**
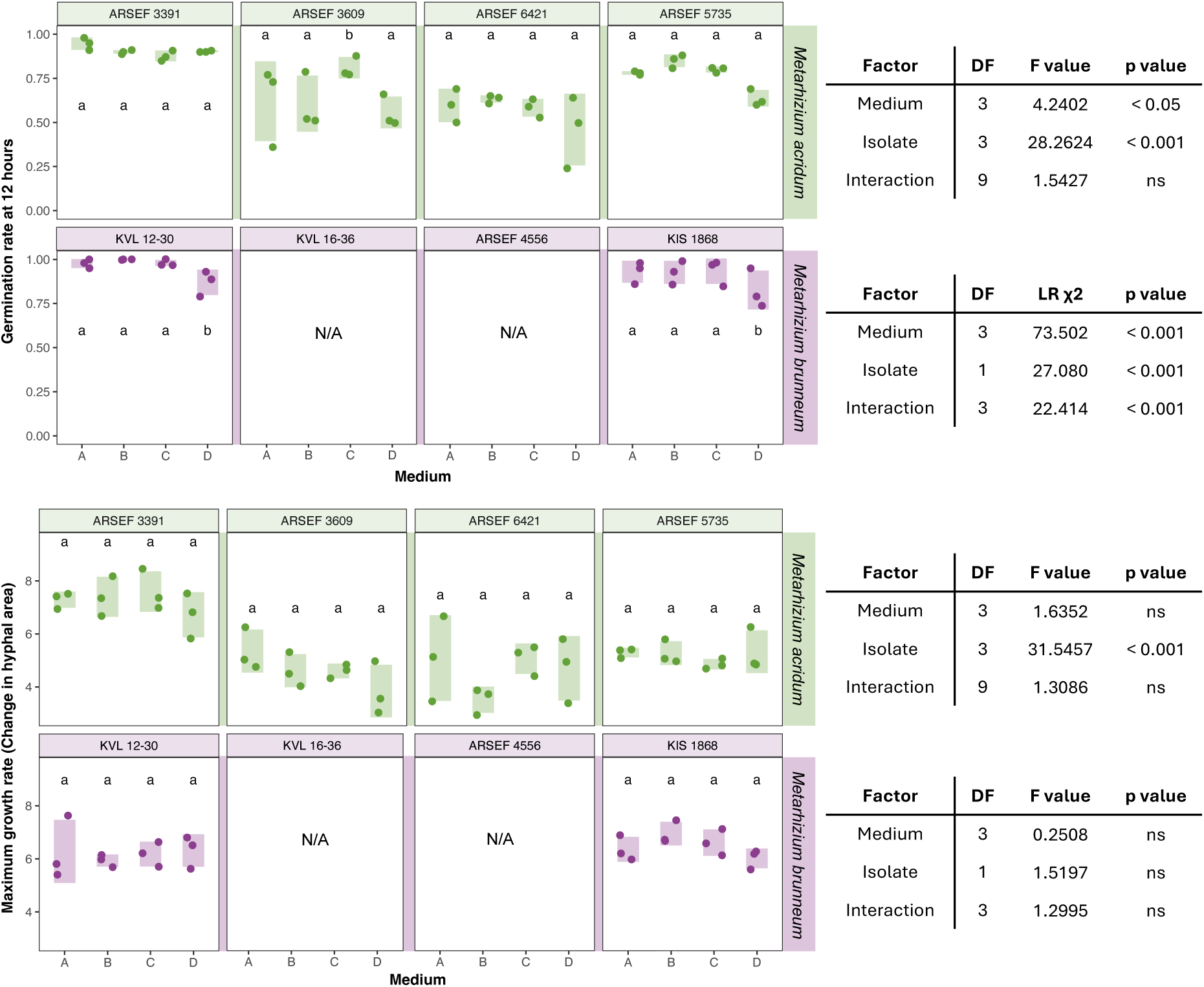
The effect of medium type origin on germination and growth in *Metarhizium acridum* and *Metarhizium brunneum*. The medium of origin had little to no effect on the germination rate at 12 hours. Exceptions included ARSEF 3609, which showed a marginally higher germination rate when spores were produced on the unfiltered locust medium, and *M. brunneum* isolates KVL1230 and KVL1939, which exhibited a marginally lower germination rate when spores were produced on the ¼ SDAY medium. The medium of origin has no effect on the maximum growth once these conidia had germinated. In all plots, the shades area behind the plotted points represents the data’s spread from minimum to maximum values.

### 4. Medium of origin significantly impacts virulence in *M. brunneum* KVL 12-30

The medium of origin had a significant impact on virulence in *M. brunneum* KVL 12-30 on *T. molitor* larvae (**Figure 5**, **Table S1 & S3**), with significantly different hazard ratios across the medium types. When using Medium A as the reference level, larvae infected with conidia derived from Medium C and Medium D had a significantly reduced risk of mortality. In particular, the hazard of death was lower in Medium C (HR = 0.39, 95% CI: 0.17 - 0.92, p < 0.05) and Medium D (HR = 0.35, CI: 0.14 - 0.87, p < 0.05), while the hazard of death was not significantly different in Medium B (HR = 1.32, 95% CI: 0.68 – 2.59, p = 0.41). The same was not true for *M. brunneum* KIS 1868, which showed consistent mortality regardless of medium type (all HR values > 0.5, all p values > 0.2).

**Figure 5:**
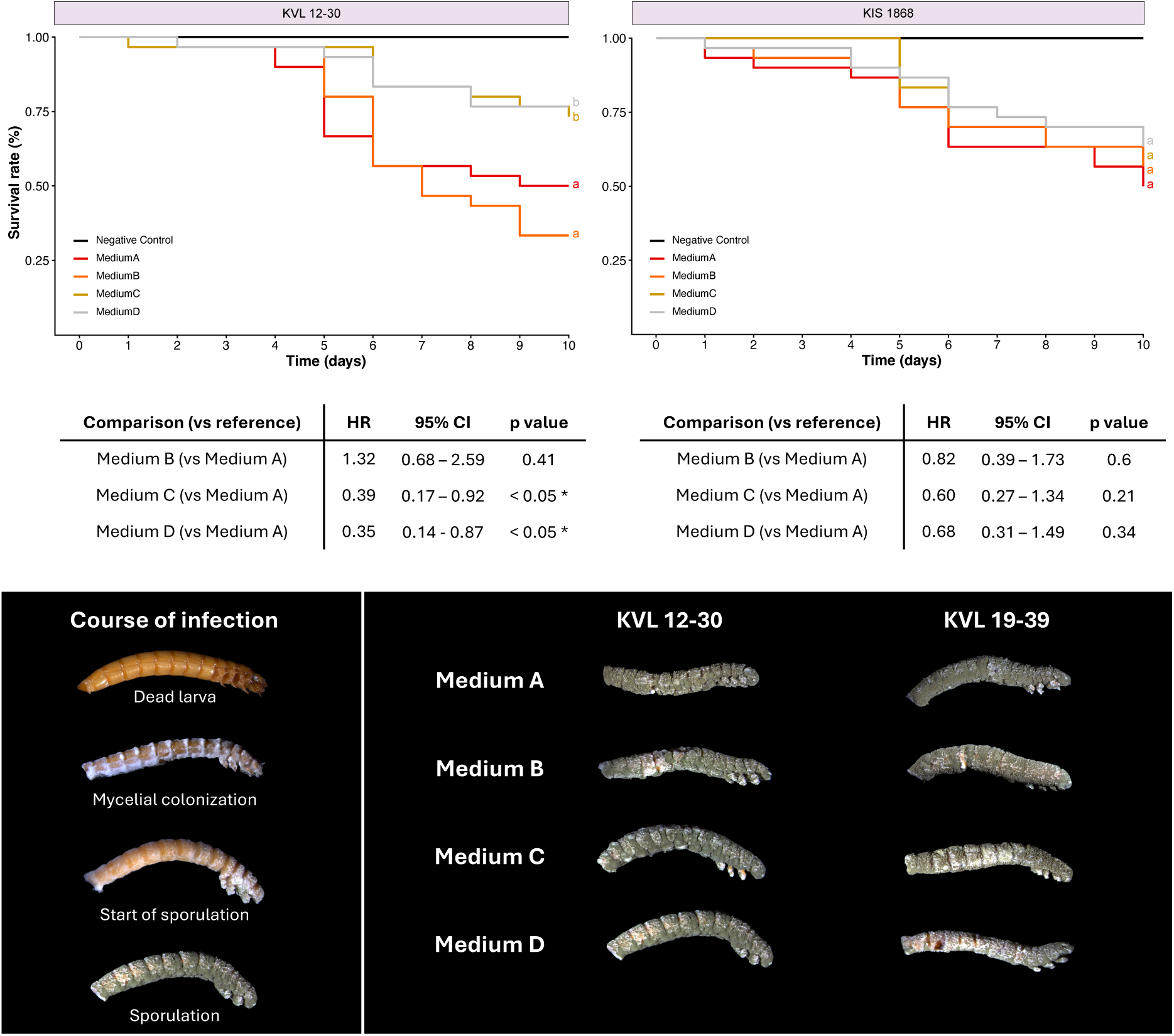
The effect of medium type origin on virulence in *Metarhizium brunneum* on *Tenebrio molitor* larvae. The medium of origin influenced the virulence of the produced conidia in *M. brunneum* KVL 12-30, with the once- and twice-filtered locust medium types exhibiting higher levels of mortality than the unfiltered locust medium and the ¼ SDAY. No differences were observed in *M. brunneum* KIS 1868. The photographs show the course of the infection, illustrating the progress from a larva on the day of recorded death, through mycelial colonization, and subsequent sporulation. Mycosis is also shown for all medium/isolate combinations four to five days after death, highlighting that spores from all four medium types were capable of infection, despite differences in virulence.

## DISCUSSION

In this study, we investigated the effect of different medium types on the phenotypic traits of two entomopathogenic fungi, *M. acridum* and *M. brunneum*. Phenotypes produced on the standard growth medium for these species, ¼ SDAY, were compared to those produced on three types of insect-derived media with the intent to determine whether the availability of host-derived nutrients and cues would impact growth, sporulation, and virulence. The presence of locust material in the growth media enhanced conidial production and decreased the average time to sporulation, while having a limited effect on radial growth. Conidia grown on these different media showed similar germination and growth rates, suggesting that medium-associated effects may be relatively short-lived. However, some differences were apparent in the virulence of the conidia produced on the different medium types, with conidia derived from two of the locust media showing higher virulence.

Conidia are produced by most fungi, from plant to insect to mammal pathogens. They have thus historically played an important role in mycological research and fungal biotechnology, being used as the inoculum for virulence, growth, stress tolerance, and host interaction assays, as well as laboratory fermentation (Butt et al., 2016; Rangel et al., 2005). In the natural infection cycles of most entomopathogens, conidia are the primary dispersal agents and mediate host attachment and subsequent germination and penetration (Quesada-Moraga et al., 2022; St. Leger, 2024). Their quantity and quality thus play an important role in the success of the pathogen. In the biocontrol context, conidia are the active ingredient of most commercial-grade fungal products (Muñiz-Paredes et al., 2017), emphasizing the importance of traits like spore yield, viability, and germination rate as key determinants of the formulation’s efficacy. It is thus essential that a deep understanding of the factors that influence spore production and quality is attained if we are to fully exploit entomopathogens as environmentally friendly biocontrol agents (Muñiz-Paredes et al., 2017).

The impact of growth medium on fungal phenotypes has been thoroughly investigated in phytopathogens (Moura et al., 2020; Overy et al., 2006), entomopathogens (Ibrahim et al., 2002; Sala et al., 2023; Vidal et al., 1998), and fungi with relevance to various biotechnology industries (Ahamed and Vermette, 2009; Laxman et al., 2005). Although studies focused on the effect of host-derived nutrients and cues in growth media are limited, this has been studied in *Penicillium* species (Overy et al., 2006) and *Metarhizium anisopliae* (Ibrahim et al., 2002). For *Penicillium*, macerated tulip bulbs were used to produce a host-derived medium that was compared to commonly used growth media. Growth on the tulip medium resulted in the production of low abundance metabolites which were also produced *in planta*, suggesting a role in the fungus-plant interaction and highlighting the promise of host-derived media as a proxy for *in vivo* experimentation (Overy et al., 2006). Similarly, when *M. anisopliae* was grown on medium containing a *Myzus persicae* homogenate, the resulting conidia germinated at a higher rate and appressorial production on host cuticles was higher, suggesting that the provision of host-derived material prior to sporulation can impact the quality of the resulting conidia (Ibrahim et al., 2002).

In the present study, locust-derived media enhanced conidial production in both *M. acridum* and *M. brunneum*. In almost all cases, the unfiltered locust medium resulted in the production of more conidia than the control medium, illustrating that host-derived materials are important for conidiation in both species. Although the precise effects were isolate-dependent, filtration of the locust-derived medium tended to impact spore production in *M. brunneum* more so than in *M. acridum*. There is thus a species-specific sensitivity to medium composition and complexity which may reflect differences in the host ranges of the two species. Together, these results indicate that incorporating host-derived material into growth substrates could be an effective strategy to optimize conidial yield in *Metarhizium* species, particularly those being developed for use in biopesticide formulations.

Clear signs of sporulation appeared earlier in the three locust media than in the control medium, as the first signs of sporulation in the control medium were not detected within the seven days of measurement. Furthermore, despite the fact that unfiltered locust medium typically resulted in the highest number of spores at the end point of the experiment, the once- and twice-filtered medium consistently resulted in shorter times to first sporulation. This may reflect some level of nutrient starvation and given that conidia are produced once the host nutrients have been fully exploited (Rangel et al., 2015; Shik et al., 2025), this may mimic the natural infection cycle. However, nutritive stress has also been shown to result in low conidial yield (Rangel et al., 2015) and thus, given additional time, mycelium grown on a richer substrate may have the nutritional capacity to produce more spores- as was seen in this study. This alludes to a trade-off between developmental speed and spore yield which has to be considered for rapid versus high-output biocontrol manufacturing strategies.

In one of the *M. brunneum* isolates (KVL 12-30), conidia produced on the two filtered locust media were more virulent than those produced on the unfiltered locus medium and the ¼ SDAY control medium. A similar phenomenon has been shown in *Conidiobolus coronatus* which, when grown on artificial medium supplemented with homogenized *Galleria mellonella* (greater wax moth) larvae or specific cuticular compounds, exhibits higher levels of virulence on *G. mellonella* (Włóka et al., 2022). It is hypothesized that the provision of insect-derived material may prime the conidia for infection, likely by activating genetic, epigenetic, and/or biochemical pathways associated with host recognition, cuticle degradation, and other pathogenicity markers. The fact that virulence was only enhanced in the two insect-derived media types that had been filtered and was not enhanced in the unfiltered medium type suggests that nutritive stress may also influence subsequent virulence. In fact, it has previously been shown that nutritive stress, in the form of growth on minimal medium, can result in the production of spores that are more virulent to insects and more stress tolerant (Rangel et al., 2015).

That the provision of host-derived nutrients and cues can enhance both sporulation and virulence is a phenomenon that can be strategically exploited to improve commercial biopesticide formulations, providing a promising avenue to increase the efficacy of entomopathogenic fungi as biocontrol agents. By incorporating insect-based materials into existing large-scale production protocols, two major hurdles may be overcome: 1) the production of conidia in much greater quantities and 2) the production of conidia that deliver more consistent in-field virulence and thus better pest management. Furthermore, if nutritive stress conditions can be mimicked within these systems, the production of more stress tolerant conidia may be possible, thereby enhancing in-field longevity and performance. Given the urgent need to transition from chemical-based pesticides to more ecologically friendly alternatives, a protocol that enhances the viability and virulence of entomopathogenic fungi using host-derived material provides a biologically informed solution, ultimately supporting more sustainable and resilient agricultural systems.

It is notable that an increase in virulence on *T. molitor* was observed for *M. brunneum* KVL 12-30 conidia produced on the locust-derived media. This suggests that generic insect-derived cues were sufficient to prime the conidia for infection, and identifying and characterizing specific insect-derived cues detected by these fungi should be a priority for future research programs. The fact that the enhancement occurred despite there being a mismatch between the insect species used in the *in vitro* medium and in the *in vivo* virulence bioassay indicates that conserved insect signals can trigger a broad-spectrum virulence response. However, the fact that virulence was not impacted by medium in *M. brunneum* KIS 1868 underscores that this phenotypic plasticity is strain dependent. Consequently, while generic priming may offer a viable strategy for some isolates and species, there may be additional host-specific cues that could further enhance virulence on a specific insect species if this species is used in the growth medium.

Germination of the conidia and their subsequent growth and virulence on *T. molitor* were used as proxies of spore quality. Medium of origin played almost no role in the germination of the *M. acridum* conidia, but *M. brunneum* conidia germinated at a higher rate when being produced on any of the three locust media compared to the control medium. Additionally, the growth of the germinated conidia was consistent, regardless of medium of origin in both species. Taken together, these results suggest that the effects of the medium of origin are short-lived for some phenotypes, like germination and growth rate; but are longer lived for others, like virulence. Therefore, while host-derived media can be used as a rapid conidia production tool, care should be taken to prevent confounding downstream experiments.

There were highly variable phenotypic responses across the different isolates of the two species. This was particularly notable in *M. brunneum*, where two of the four isolates could not be used in the medium of origin experiments as insufficient conidia were produced on the control medium despite also growing fastest on this medium. The diversity observed in *M. brunneum* may reflect its generalist ecology and associated metabolic flexibility, which may allow certain isolates to perform well across diverse substrates, while others depend on specific environmental cues for growth and sporulation. It may also be a reflection of the genetic diversity that is known to exist in *M. brunneum*. More importantly, the intra- and inter-species differences observed here highlight that biocontrol interventions cannot rely on uniform cultivation protocols. Instead, the effective development and deployment of fungal biopesticides requires isolate-specific optimization to ensure reliable conidial production and consistent performance in field.

Passaging pathogenic fungi through their hosts- whether insects or plants- is a common practice in mycology. It is typically done to reinvigorate a fungal isolate that has degenerated after successive subculturing on artificial media (Butt et al., 2006). Exposure to its natural host can restore key phenotypic traits, including culture characteristics and virulence. This is likely due to the reactivation of pathways associated with host interaction. However, there are significant difficulties with passaging, given that a sterile environment cannot be maintained and thus isolating the fungus of interest can be experimentally challenging. The development of a host-derived medium may circumvent the need for passaging by exposing the fungus to cues and signals from the host in an axenic environment. The results here show that that growth on host-derived media enhances fungal vigour, particularly in terms of conidial production and growth, suggesting that such a medium may recapitulate the benefits of traditional passaging. As such, this approach represents a promising methodology for maintaining entomopathogen phenotypic efficacy during industrial fermentation.

## LIST OF ABBREVIATIONS

ANOVA: Analysis of variance
ARSEF: Agricultural Research Service Collection of Entomopathogenic Fungi
CI: Confidence intervals
coxph: cox proportional hazard
DF: Degrees of freedom
FDR: False discovery rate
HR: Hazard ratio
KVL: Kongelige Veterinær- og Landbohøjskole *(English: Royal Veterinary and Agricultural)*
KIS: Kmetijski inštitut Slovenije *(English: Agricultural Institute of Slovenia)*
LR: Likelihood ration
SDY: Sabouraud dextrose with yeast extract
SDAY: Sabouraud dextrose with yeast extract agar

## DECLARATIONS

## 1. Ethical approval and consent to participate

Not applicable

## 2. Consent for publication

Not applicable

## 3. Availability of data and materials

All data generated and analysed during this study are included in the supplementary information files, including Tables S1 – S3.

## 6. Authors’ contributions

AMW: Conceptualization; data curation; formal analysis; funding acquisition; investigation; methodology; project administration; visualization; writing—original draft; writing—review and editing.

HHDFL: Conceptualization; funding acquisition; investigation; supervision; writing—review and editing.

## 7. Acknowledgements

The authors would like to thank Dr Jaka Razinger for sharing *Metarhizium brunneum* isolate KIS 1868 for use in this study.

