## Supplementary material for "Improving the production and virulence of entomopathogenic fungi for biological control using insect-derived *in vitro* culture medium": Supp Tables

**Table S1:** Summary of all models and statistics

| Spore Count (Solid Medium) | | | | | | | |
|---|---|---|---|---|---|---|---|
| ***M. acridum*** | **Distribution** | **Model** | **Anova** | **Anova results** | | | |
| | Negative Binomial | glm.nb, AverageSporeCount ~ Medium * Isolate | Type II | Factor | Degrees of Freedom | LR χ2 | P value |
| | | | | Medium | 3 | 145.42 | p < 0.001 |
| | | | | Isolate | 3 | 142.78 | p < 0.001 |
| | | | | Medium:Isolate | 9 | 197.64 | p < 0.001 |
| ***M. brunneum*** | **Distribution** | **Model** | **Anova** | **Anova results** | | | |
| | Negative Binomial | glm.nb, AverageSporeCount ~ Medium * Isolate | Type II | Factor | Degrees of Freedom | LR χ2 | P value |
| | | | | Medium | 3 | 212.57 | p < 0.001 |
| | | | | Isolate | 3 | 147.55 | p < 0.001 |
| | | | | Medium:Isolate | 9 | 934.71 | p < 0.001 |

| Time to Sporulation (Solid Medium) | | | | | | | |
|---|---|---|---|---|---|---|---|
| ***M. acridum*** | **Distribution** | **Model** | **Anova** | **Anova results** | | | |
| | Gamma | glm, HourToFirstSporulation ~ Medium + Isolate | Type II | Factor | Degrees of Freedom | LR χ2 | P value |
| | | Family = gamma(link = "inverse") | | Medium | 2 | 306.81 | p < 0.001 |
| | | | | Isolate | 3 | 1035.18 | p < 0.001 |
| | | | | Medium:Isolate | 6 | 32.22 | p < 0.001 |
| ***M. brunneum*** | **Distribution** | **Model** | **Anova** | **Anova results** | | | |
| | Gamma | glm, HourToFirstSporulation ~ Medium + Isolate | Type II | Factor | Degrees of Freedom | LR χ2 | P value |
| | | Family = gamma(link = "inverse") | | Medium | 2 | 130.943 | p < 0.001 |
| | | | | Isolate | 2 | 108.321 | p < 0.001 |
| | | | | Medium:Isolate | 4 | 2.238 | ns |

| Radial Growth Rate (Solid Medium) | | | | |
|---|---|---|---|---|
| ***M. acridum*** | **Distribution** | **Model** | **Anova** | **Anova results** |

| | | | | | | | |
|---|---|---|---|---|---|---|---|
| | Gaussian | lm, GrowthRate ~ Medium * Isolate | Type II | Factor | Degrees of Freedom | F value | P value |
| | | | | Medium | 3 | 39.9905 | p < 0.001 |
| | | | | Isolate | 3 | 14.9252 | p < 0.001 |
| | | | | Medium:Isolate | 9 | 4.0061 | p < 0.01 |
| | | | | Residuals | 32 | | |
| ***M. brunneum*** | **Distribution** | **Model** | **Anova** | **Anova results** | | | |
| | Gaussian | lm, GrowthRate ~ Medium * Isolate | Type II | Factor | Degrees of Freedom | F value | P value |
| | | | | Medium | 3 | 31.221 | p < 0.001 |
| | | | | Isolate | 3 | 31.697 | p < 0.001 |
| | | | | Medium:Isolate | 9 | 12.169 | p < 0.001 |
| | | | | Residuals | 32 | | |

| **Germination at 12 hours (LiquidMedium)** | | | | | | | |
|---|---|---|---|---|---|---|---|
| ***M. acridum*** | **Distribution** | **Model** | **Anova** | **Anova results** | | | |
| | Gaussian | lm, GrowthRate ~ Medium * Isolate | Type II | Factor | Degrees of Freedom | F value | P value |
| | | | | Medium | 3 | 4.2402 | p < 0.05 |
| | | | | Isolate | 3 | 28.2624 | p < 0.001 |
| | | | | Medium:Isolate | 9 | 1.5427 | ns |
| | | | | Residuals | 32 | | |
| ***M. brunneum*** | **Distribution** | **Model** | **Anova** | **Anova results** | | | |
| | Binomial | glm, GermRate_12hr ~ Medium * Isolate | Type II | Factor | Degrees of Freedom | F value | P value |
| | | Family = binomial, Weights = 100 | | Medium | 3 | 73.502 | p < 0.001 |
| | | | | Isolate | 1 | 27.08 | p < 0.001 |
| | | | | Medium:Isolate | 3 | 22.414 | p < 0.001 |

| **Maximum Growth Rate (LiquidMedium)** | | | | | | | |
|---|---|---|---|---|---|---|---|
| ***M. acridum*** | **Distribution** | **Model** | **Anova** | **Anova results** | | | |
| | Gaussian | lm, GrowthRate ~ Medium * Isolate | Type II | Factor | Degrees of Freedom | F value | P value |

| | | | | | | | |
|---|---|---|---|---|---|---|---|
| | | | | Medium | 3 | 1.6352 | ns |
| | | | | Isolate | 3 | 31.5457 | p < 0.001 |
| | | | | Medium:Isolate | 9 | 1.3086 | ns |
| | | | | Residuals | 32 | | |
| ***M. brunneum*** | **Distribution** | **Model** | **Anova** | **Anova results** | | | |
| | Gaussian | lm, GrowthRate ~ Medium * Isolate | Type II | Factor | Degrees of Freedom | F value | P value |
| | | | | Medium | 3 | 0.2508 | ns |
| | | | | Isolate | 1 | 1.5197 | ns |
| | | | | Medium:Isolate | 3 | 1.2995 | ns |
| | | | | Residuals | 16 | | |

| **Virulence in *Tenebrio molitor*** | | | | | | | |
|---|---|---|---|---|---|---|---|
| ***M. brunneum*** | **Model** | | | | | | |
| | coxph(Surv(Survival_Time, Censorship) ~ Medium + Repetition) | | | | | | |
| **KVL 12-30** | n = 120, number of events = 50 | | | | | | |
| | **MediumMediumB** | 0.2793 | 1.3221 | 0.3426 | 0.815 | 0.4149 | |
| | **MediumMediumC** | -0.9386 | 0.3912 | 0.4385 | -2.141 | 0.0323 | * |
| | **MediumMediumD** | -1.0415 | 0.3529 | 0.4583 | -2.272 | 0.0231 | * |
| | **Repetition2** | -0.1422 | 0.8674 | 0.3385 | -0.42 | 0.6744 | |
| | **Repetition3** | -0.2735 | 0.7607 | 0.3504 | -0.78 | 0.4351 | |
| Concordance = 0.658 (se = 0.036) | | | | | | | |
| Likelihood ratio test = 16.18 on 5 df, p = 0.006 | | | | | | | |
| Wald test = 14.88 on 5 df, p = 0.01 | | | | | | | |
| Score (logrank) test = 16.54 on 5 df, p = 0.005 | | | | | | | |
| | **chisq** | **df** | **p** | | | | |
| **Medium** | 5.98 | 3 | 0.11 | | | | |

| | | | | | | | | |
|---|---|---|---|---|---|---|---|---|
| **Repetition** | 0.564 | 2 | 0.75 | | | | | |
| **GLOBAL** | 6.7 | 5 | 0.24 | | | | | |
| **KVL 19-30** | n = 120, number of events = 49 | | | | | | | |
| | | | **coef** | **exp(coef)** | **se(coef)** | **z** | **Pr(>\|z\|)** | |
| | **MediumMediumB** | | -0.1966 | 0.8215 | 0.3793 | -0.518 | 0.6042 | |
| | **MediumMediumC** | | -0.5086 | 0.6014 | 0.4086 | -1.244 | 0.2133 | |
| | **MediumMediumD** | | -0.3811 | 0.6831 | 0.398 | -0.958 | 0.3383 | |
| | **Repetition2** | | -0.6262 | 0.5346 | 0.3543 | -1.767 | 0.0772 | . |
| | **Repetition3** | | -0.4138 | 0.6611 | 0.3392 | -1.22 | 0.2226 | |
| Concordance = 0.613 (se = 0.038 ) | | | | | | | | |
| Likelihood ratio test = 5.55 on 5 df, p = 0.4 | | | | | | | | |
| Wald test = 5.6 on 5 df, p = 0.3 | | | | | | | | |
| Score (logrank) test = 5.74 on 5 df, p = 0.3 | | | | | | | | |
| | **chisq** | **df** | **p** | | | | | |
| **Medium** | 0.242 | 3 | 0.97 | | | | | |
| **Repetition** | 0.648 | 2 | 0.72 | | | | | |
| **GLOBAL** | 0.865 | 5 | 0.97 | | | | | |

**Table S2:** All phenotypic measurements regarding sporulation, growth, and germination

| Plate | Well | Isolate | Species | Medium | Rep | FirstSporulationImage | HourToFirstSporulation | AverageAt31 | AverageAt61 | GrowthRate1 |
|---|---|---|---|---|---|---|---|---|---|---|
| A1 | A1 | ARSEF3391 | M_acridum | A | 1 | 45 | 88 | 0.847 | 1.2175 | 0.3705 |
| A1 | B1 | ARSEF3391 | M_acridum | A | 2 | 45 | 88 | 0.9475 | 1.245 | 0.2975 |
| A1 | C1 | ARSEF3391 | M_acridum | A | 3 | 45 | 88 | 0.812 | 1.2795 | 0.4675 |
| A1 | A2 | ARSEF3609 | M_acridum | A | 1 | 55 | 108 | 0.758 | 1.266 | 0.508 |
| A1 | B2 | ARSEF3609 | M_acridum | A | 2 | 54 | 106 | 0.826 | 1.288 | 0.462 |
| A1 | C2 | ARSEF3609 | M_acridum | A | 3 | 59 | 116 | 0.786 | 1.269 | 0.483 |
| A1 | A3 | ARSEF6421 | M_acridum | A | 1 | 48 | 94 | 0.7965 | 1.198 | 0.4015 |
| A1 | B3 | ARSEF6421 | M_acridum | A | 2 | 52 | 102 | 0.8485 | 1.2005 | 0.352 |
| A1 | C3 | ARSEF6421 | M_acridum | A | 3 | 48 | 94 | 0.8085 | 1.2155 | 0.407 |
| A1 | A4 | ARSEF5735 | M_acridum | A | 1 | 40 | 78 | 0.6985 | 1.062 | 0.3635 |
| A1 | B4 | ARSEF5735 | M_acridum | A | 2 | 40 | 78 | 0.6755 | 1.153 | 0.4775 |
| A1 | C4 | ARSEF5735 | M_acridum | A | 3 | 40 | 78 | 0.616 | 1.0705 | 0.4545 |
| A2 | A3 | ARSEF4556 | M_brunneum | A | 1 | NA | NA | 1.0425 | 1.427 | 0.3845 |
| A2 | B3 | ARSEF4556 | M_brunneum | A | 2 | NA | NA | 1.076 | 1.4445 | 0.3685 |
| A2 | C3 | ARSEF4556 | M_brunneum | A | 3 | NA | NA | 0.9835 | 1.417 | 0.4335 |
| A2 | A1 | KVL1230 | M_brunneum | A | 1 | 45 | 88 | 0.898 | 1.4655 | 0.5675 |
| A2 | B1 | KVL1230 | M_brunneum | A | 2 | 45 | 88 | 0.8665 | 1.462 | 0.5955 |
| A2 | C1 | KVL1230 | M_brunneum | A | 3 | 45 | 88 | 0.842 | 1.4445 | 0.6025 |
| A2 | A2 | KVL1636 | M_brunneum | A | 1 | 44 | 86 | 1.006 | 1.57 | 0.564 |
| A2 | B2 | KVL1636 | M_brunneum | A | 2 | 44 | 86 | 0.9695 | 1.615 | 0.6455 |
| A2 | C2 | KVL1636 | M_brunneum | A | 3 | 44 | 86 | 0.952 | 1.587 | 0.635 |
| A2 | A4 | KVL1939 | M_brunneum | A | 1 | 40 | 78 | 0.8995 | 1.448 | 0.5485 |
| A2 | B4 | KVL1939 | M_brunneum | A | 2 | 39 | 76 | 0.919 | 1.462 | 0.543 |
| A2 | C4 | KVL1939 | M_brunneum | A | 3 | 39 | 76 | 0.874 | 1.403 | 0.529 |
| B1 | A1 | ARSEF3391 | M_acridum | B | 1 | 49 | 96 | 0.6925 | 1.177 | 0.4845 |
| B1 | B1 | ARSEF3391 | M_acridum | B | 2 | 49 | 96 | 0.717 | 1.2395 | 0.5225 |

| | | | | | | | | | | |
|---|---|---|---|---|---|---|---|---|---|---|
| B1 | C1 | ARSEF3391 | M_acridum | B | 3 | 48 | 94 | 0.6785 | 1.2045 | 0.526 |
| B1 | A2 | ARSEF3609 | M_acridum | B | 1 | 59 | 116 | 0.731 | 1.3055 | 0.5745 |
| B1 | B2 | ARSEF3609 | M_acridum | B | 2 | 59 | 116 | 0.919 | 1.455 | 0.536 |
| B1 | C2 | ARSEF3609 | M_acridum | B | 3 | 59 | 116 | 0.7765 | 1.375 | 0.5985 |
| B1 | A3 | ARSEF6421 | M_acridum | B | 1 | 53 | 104 | 0.752 | 1.222 | 0.47 |
| B1 | B3 | ARSEF6421 | M_acridum | B | 2 | 54 | 106 | 0.9155 | 1.3665 | 0.451 |
| B1 | C3 | ARSEF6421 | M_acridum | B | 3 | 53 | 104 | 0.794 | 1.2965 | 0.5025 |
| B1 | A4 | ARSEF5735 | M_acridum | B | 1 | 45 | 88 | 0.668 | 1.135 | 0.467 |
| B1 | B4 | ARSEF5735 | M_acridum | B | 2 | 45 | 88 | 0.769 | 1.161 | 0.392 |
| B1 | C4 | ARSEF5735 | M_acridum | B | 3 | 45 | 88 | 0.6685 | 1.154 | 0.4855 |
| B2 | A3 | ARSEF4556 | M_brunneum | B | 1 | NA | NA | 1.0685 | 1.5595 | 0.491 |
| B2 | B3 | ARSEF4556 | M_brunneum | B | 2 | NA | NA | 1.2185 | 1.5945 | 0.376 |
| B2 | C3 | ARSEF4556 | M_brunneum | B | 3 | NA | NA | 1.107 | 1.5595 | 0.4525 |
| B2 | A1 | KVL1230 | M_brunneum | B | 1 | 46 | 90 | 0.9505 | 1.5075 | 0.557 |
| B2 | B1 | KVL1230 | M_brunneum | B | 2 | 47 | 92 | 1.013 | 1.5735 | 0.5605 |
| B2 | C1 | KVL1230 | M_brunneum | B | 3 | 46 | 90 | 0.968 | 1.5145 | 0.5465 |
| B2 | A2 | KVL1636 | M_brunneum | B | 1 | 46 | 90 | 1.053 | 1.709 | 0.656 |
| B2 | B2 | KVL1636 | M_brunneum | B | 2 | 47 | 92 | 1.217 | 1.8275 | 0.6105 |
| B2 | C2 | KVL1636 | M_brunneum | B | 3 | 46 | 90 | 1.109 | 1.8275 | 0.7185 |
| B2 | A4 | KVL1939 | M_brunneum | B | 1 | 41 | 80 | 0.9175 | 1.4725 | 0.555 |
| B2 | B4 | KVL1939 | M_brunneum | B | 2 | 41 | 80 | 0.9535 | 1.5125 | 0.559 |
| B2 | C4 | KVL1939 | M_brunneum | B | 3 | 41 | 80 | 0.95 | 1.5595 | 0.6095 |
| C1 | A1 | ARSEF3391 | M_acridum | C | 1 | 54 | 106 | 0.717 | 1.274 | 0.557 |
| C1 | B1 | ARSEF3391 | M_acridum | C | 2 | 54 | 106 | 0.7695 | 1.33 | 0.5605 |
| C1 | C1 | ARSEF3391 | M_acridum | C | 3 | 54 | 106 | 0.6685 | 1.288 | 0.6195 |
| C1 | A2 | ARSEF3609 | M_acridum | C | 1 | 63 | 124 | 0.752 | 1.427 | 0.675 |
| C1 | B2 | ARSEF3609 | M_acridum | C | 2 | 62 | 122 | 0.9155 | 1.5905 | 0.675 |
| C1 | C2 | ARSEF3609 | M_acridum | C | 3 | 63 | 124 | 0.8045 | 1.4725 | 0.668 |
| C1 | A3 | ARSEF6421 | M_acridum | C | 1 | 56 | 110 | 0.7695 | 1.3665 | 0.597 |

| | | | | | | | | | | |
|---|---|---|---|---|---|---|---|---|---|---|
| C1 | B3 | ARSEF6421 | M_acridum | C | 2 | 57 | 112 | 0.8875 | 1.443 | 0.5555 |
| C1 | C3 | ARSEF6421 | M_acridum | C | 3 | 56 | 110 | 0.839 | 1.4605 | 0.6215 |
| C1 | A4 | ARSEF5735 | M_acridum | C | 1 | 47 | 92 | 0.6825 | 1.246 | 0.5635 |
| C1 | B4 | ARSEF5735 | M_acridum | C | 2 | 47 | 92 | 0.804 | 1.396 | 0.592 |
| C1 | C4 | ARSEF5735 | M_acridum | C | 3 | 47 | 92 | 0.7345 | 1.396 | 0.6615 |
| C2 | A3 | ARSEF4556 | M_brunneum | C | 1 | 72 | 142 | 1.041 | 1.6605 | 0.6195 |
| C2 | B3 | ARSEF4556 | M_brunneum | C | 2 | 65 | 128 | 1.26 | 1.758 | 0.498 |
| C2 | C3 | ARSEF4556 | M_brunneum | C | 3 | 79 | 156 | 1.1105 | 1.671 | 0.5605 |
| C2 | A1 | KVL1230 | M_brunneum | C | 1 | 53 | 104 | 0.947 | 1.587 | 0.64 |
| C2 | B1 | KVL1230 | M_brunneum | C | 2 | 53 | 104 | 0.9505 | 1.5285 | 0.578 |
| C2 | C1 | KVL1230 | M_brunneum | C | 3 | 53 | 104 | 0.9505 | 1.577 | 0.6265 |
| C2 | A2 | KVL1636 | M_brunneum | C | 1 | 51 | 100 | 1.0755 | 1.7475 | 0.672 |
| C2 | B2 | KVL1636 | M_brunneum | C | 2 | 60 | 118 | 1.215 | 1.8555 | 0.6405 |
| C2 | C2 | KVL1636 | M_brunneum | C | 3 | 51 | 100 | 1.0545 | 1.775 | 0.7205 |
| C2 | A4 | KVL1939 | M_brunneum | C | 1 | 45 | 88 | 0.94 | 1.5315 | 0.5915 |
| C2 | B4 | KVL1939 | M_brunneum | C | 2 | 45 | 88 | 1.006 | 1.6115 | 0.6055 |
| C2 | C4 | KVL1939 | M_brunneum | C | 3 | 45 | 88 | 0.9605 | 1.57 | 0.6095 |
| D1 | A1 | ARSEF3391 | M_acridum | D | 1 | NA | NA | 0.7885 | 1.3785 | 0.59 |
| D1 | B1 | ARSEF3391 | M_acridum | D | 2 | NA | NA | 0.813 | 1.413 | 0.6 |
| D1 | C1 | ARSEF3391 | M_acridum | D | 3 | NA | NA | 0.7435 | 1.382 | 0.6385 |
| D1 | A2 | ARSEF3609 | M_acridum | D | 1 | NA | NA | 0.7975 | 1.344 | 0.5465 |
| D1 | B2 | ARSEF3609 | M_acridum | D | 2 | NA | NA | 0.656 | 1.302 | 0.646 |
| D1 | C2 | ARSEF3609 | M_acridum | D | 3 | NA | NA | 0.7485 | 1.26 | 0.5115 |
| D1 | A3 | ARSEF6421 | M_acridum | D | 1 | 58 | 114 | 0.8615 | 1.288 | 0.4265 |
| D1 | B3 | ARSEF6421 | M_acridum | D | 2 | 58 | 114 | 0.823 | 1.2495 | 0.4265 |
| D1 | C3 | ARSEF6421 | M_acridum | D | 3 | 58 | 114 | 0.8405 | 1.302 | 0.4615 |
| D1 | A4 | ARSEF5735 | M_acridum | D | 1 | 58 | 114 | 0.7655 | 1.1805 | 0.415 |
| D1 | B4 | ARSEF5735 | M_acridum | D | 2 | 58 | 114 | 0.785 | 1.142 | 0.357 |
| D1 | C4 | ARSEF5735 | M_acridum | D | 3 | 58 | 114 | 0.698 | 1.14 | 0.442 |

| | | | | | | | | | |
|---|---|---|---|---|---|---|---|---|---|
| D2 | A3 | ARSEF4556 | M_brunneum | D | 1 | NA | NA | 1.034 | 1.789 | 0.755 |
| D2 | B3 | ARSEF4556 | M_brunneum | D | 2 | NA | NA | 1.0425 | 1.838 | 0.7955 |
| D2 | C3 | ARSEF4556 | M_brunneum | D | 3 | NA | NA | 1.0355 | 1.786 | 0.7505 |
| D2 | A1 | KVL1230 | M_brunneum | D | 1 | NA | NA | 0.8115 | 1.417 | 0.6055 |
| D2 | B1 | KVL1230 | M_brunneum | D | 2 | NA | NA | 0.842 | 1.4915 | 0.6495 |
| D2 | C1 | KVL1230 | M_brunneum | D | 3 | NA | NA | 0.822 | 1.462 | 0.64 |
| D2 | A2 | KVL1636 | M_brunneum | D | 1 | NA | NA | 0.9785 | 1.789 | 0.8105 |
| D2 | B2 | KVL1636 | M_brunneum | D | 2 | NA | NA | 0.9785 | 1.8345 | 0.856 |
| D2 | C2 | KVL1636 | M_brunneum | D | 3 | NA | NA | 0.9505 | 1.65 | 0.6995 |
| D2 | A4 | KVL1939 | M_brunneum | D | 1 | NA | NA | 0.877 | 1.392 | 0.515 |
| D2 | B4 | KVL1939 | M_brunneum | D | 2 | NA | NA | 0.888 | 1.42 | 0.532 |
| D2 | C4 | KVL1939 | M_brunneum | D | 3 | NA | NA | 0.8775 | 1.403 | 0.5255 |

[redacted]

| GrowthRate2 | AverageSporeCount | MaxSpores | MinSpore_PostMax | Time_Max | Time_Min | Perc50 | TimeTo50perc | GermRate_12hr | GermRate_24hr |
|---|---|---|---|---|---|---|---|---|---|
| 0.1482 | 3.88E+07 | 38 | 0 | 0 | 20 | 19 | 7 | 0.95 | 1.00 |
| 0.119 | 4.61E+07 | 54 | 0 | 1 | 21 | 27 | 6 | 0.91 | 0.98 |
| 0.187 | 3.99E+07 | 40 | 0 | 0 | 18 | 20 | 4 | 0.98 | 1.00 |
| 0.2032 | 4.37E+07 | 64 | 0 | 4 | 17 | 32 | 10 | 0.73 | 1.00 |
| 0.1848 | 5.30E+07 | 77 | 0 | 0 | 19 | 38.5 | 13 | 0.36 | 1.00 |
| 0.1932 | 3.96E+07 | 69 | 0 | 6 | 18 | 34.5 | 10 | 0.77 | 0.97 |
| 0.1606 | 4.90E+07 | 78 | 0 | 3 | 22 | 39 | 12 | 0.50 | 0.96 |
| 0.1408 | 6.27E+07 | 93 | 0 | 2 | 21 | 46.5 | 11 | 0.69 | 1.00 |
| 0.1628 | 5.81E+07 | 98 | 0 | 0 | 23 | 49 | 12 | 0.60 | 0.99 |
| 0.1454 | 6.56E+07 | 79 | 0 | 0 | 24 | 39.5 | 11 | 0.78 | 0.99 |
| 0.191 | 6.40E+07 | 94 | 0 | 4 | 23 | 47 | 9 | 0.77 | 0.98 |
| 0.1818 | 7.44E+07 | 92 | 0 | 0 | 24 | 46 | 10 | 0.79 | 0.97 |
| 0.1538 | 1.07E+07 | 40 | 0 | 0 | 13 | 20 | 5 | 0.98 | 1.00 |
| 0.1474 | 8.03E+06 | 37 | 0 | 0 | 17 | 18.5 | 3 | 0.97 | 1.00 |
| 0.1734 | 1.15E+07 | 35 | 0 | 0 | 10 | 17.5 | 3 | 1.00 | 1.00 |
| 0.227 | 3.72E+07 | 58 | 0 | 0 | 17 | 29 | 6 | 0.95 | 1.00 |
| 0.2382 | 3.70E+07 | 46 | 0 | 0 | 11 | 23 | 6 | 1.00 | 1.00 |
| 0.241 | 6.24E+07 | 53 | 0 | 1 | 12 | 26.5 | 6 | 0.98 | 0.98 |
| 0.2256 | 4.42E+07 | 57 | 0 | 1 | 17 | 28.5 | 6 | 0.96 | 1.00 |
| 0.2582 | 4.18E+07 | 46 | 0 | 0 | 16 | 23 | 6 | 0.98 | 1.00 |
| 0.254 | 3.43E+07 | 42 | 0 | 0 | 16 | 21 | 7 | 0.93 | 1.00 |
| 0.2194 | 3.43E+07 | 61 | 0 | 2 | 12 | 30.5 | 6 | 0.98 | 0.98 |
| 0.2172 | 5.22E+07 | 61 | 0 | 2 | 17 | 30.5 | 7 | 0.95 | 0.98 |
| 0.2116 | 5.46E+07 | 88 | 0 | 3 | 18 | 44 | 7 | 0.86 | 1.00 |
| 0.1938 | 8.41E+07 | 54 | 0 | 1 | 16 | 27 | 6 | 0.91 | 1.00 |
| 0.209 | 8.97E+07 | 63 | 0 | 2 | 19 | 31.5 | 5 | 0.90 | 1.00 |

| | | | | | | | | | |
|---|---|---|---|---|---|---|---|---|---|
| 0.2104 | 6.48E+07 | 64 | 0 | 2 | 24 | 32 | 5 | 0.89 | 1.00 |
| 0.2298 | 4.37E+07 | 73 | 0 | 3 | 23 | 36.5 | 12 | 0.51 | 1.00 |
| 0.2144 | 6.43E+07 | 75 | 0 | 5 | 18 | 37.5 | 11 | 0.52 | 0.97 |
| 0.2394 | 3.91E+07 | 62 | 0 | 2 | 23 | 31 | 10 | 0.79 | 0.97 |
| 0.188 | 6.16E+07 | 77 | 0 | 1 | 20 | 38.5 | 11 | 0.64 | 0.97 |
| 0.1804 | 8.54E+07 | 77 | 0 | 5 | 22 | 38.5 | 11 | 0.65 | 1.00 |
| 0.201 | 6.59E+07 | 67 | 0 | 1 | 23 | 33.5 | 12 | 0.61 | 1.00 |
| 0.1868 | 8.03E+07 | 68 | 0 | 0 | 19 | 34 | 9 | 0.88 | 0.99 |
| 0.1568 | 8.17E+07 | 85 | 0 | 0 | 21 | 42.5 | 9 | 0.86 | 0.96 |
| 0.1942 | 7.07E+07 | 78 | 0 | 0 | 22 | 39 | 8 | 0.81 | 0.97 |
| 0.1964 | 1.39E+07 | 96 | 0 | 0 | 19 | 48 | 3 | 0.99 | 1.00 |
| 0.1504 | 2.65E+07 | 102 | 0 | 0 | 11 | 51 | 4 | 1.00 | 1.00 |
| 0.181 | 1.85E+07 | 92 | 0 | 1 | 22 | 46 | 5 | 0.93 | 1.00 |
| 0.2228 | 8.81E+07 | 54 | 0 | 3 | 12 | 27 | 6 | 1.00 | 1.00 |
| 0.2242 | 7.58E+07 | 62 | 0 | 1 | 12 | 31 | 5 | 1.00 | 1.00 |
| 0.2186 | 9.24E+07 | 48 | 0 | 0 | 12 | 24 | 6 | 1.00 | 1.00 |
| 0.2624 | 8.19E+07 | 66 | 0 | 0 | 12 | 33 | 5 | 1.00 | 1.00 |
| 0.2442 | 1.01E+08 | 65 | 0 | 0 | 11 | 32.5 | 6 | 1.00 | 1.00 |
| 0.2874 | 1.03E+08 | 62 | 0 | 1 | 12 | 31 | 5 | 1.00 | 1.00 |
| 0.222 | 5.06E+06 | 110 | 0 | 2 | 23 | 55 | 6 | 0.99 | 1.00 |
| 0.2236 | 7.71E+07 | 73 | 0 | 3 | 19 | 36.5 | 7 | 0.93 | 1.00 |
| 0.2438 | 1.01E+08 | 86 | 0 | 1 | 21 | 43 | 7 | 0.86 | 0.97 |
| 0.2228 | 6.88E+07 | 62 | 0 | 1 | 20 | 31 | 6 | 0.85 | 1.00 |
| 0.2242 | 9.19E+07 | 55 | 0 | 1 | 20 | 27.5 | 6 | 0.91 | 1.00 |
| 0.2478 | 7.18E+07 | 68 | 0 | 0 | 21 | 34 | 7 | 0.87 | 0.99 |
| 0.27 | 4.31E+07 | 72 | 0 | 2 | 21 | 36 | 11 | 0.78 | 0.96 |
| 0.27 | 6.72E+07 | 89 | 0 | 3 | 23 | 44.5 | 10 | 0.88 | 1.00 |
| 0.2672 | 3.56E+07 | 77 | 0 | 1 | 20 | 38.5 | 11 | 0.77 | 0.99 |
| 0.2388 | 6.78E+07 | 56 | 0 | 0 | 21 | 28 | 11 | 0.59 | 0.98 |

| | | | | | | | | | |
|---|---|---|---|---|---|---|---|---|---|
| 0.2222 | 7.85E+07 | 64 | 0 | 5 | 24 | 32 | 12 | 0.53 | 0.95 |
| 0.2486 | 1.51E+08 | 78 | 0 | 2 | 23 | 39 | 11 | 0.63 | 0.91 |
| 0.2254 | 1.15E+08 | 58 | 0 | 0 | 16 | 29 | 8 | 0.81 | 1.00 |
| 0.2368 | 1.32E+08 | 64 | 0 | 0 | 24 | 32 | 8 | 0.81 | 0.95 |
| 0.2646 | 1.10E+08 | 67 | 0 | 0 | 22 | 33.5 | 9 | 0.78 | 0.96 |
| 0.2478 | 3.03E+07 | 52 | 0 | 0 | 15 | 26 | 3 | 0.98 | 1.00 |
| 0.1992 | 2.70E+06 | 65 | 0 | 0 | 11 | 32.5 | 3 | 1.00 | 1.00 |
| 0.2242 | 2.04E+07 | 48 | 0 | 0 | 22 | 24 | 4 | 0.98 | 1.00 |
| 0.256 | 8.70E+07 | 63 | 0 | 1 | 19 | 31.5 | 5 | 0.97 | 1.00 |
| 0.2312 | 8.94E+07 | 61 | 0 | 0 | 15 | 30.5 | 7 | 0.97 | 0.98 |
| 0.2506 | 9.94E+07 | 45 | 0 | 2 | 9 | 22.5 | 6 | 1.00 | 1.00 |
| 0.2688 | 1.78E+08 | 41 | 0 | 0 | 10 | 20.5 | 6 | 1.00 | 1.00 |
| 0.2562 | 1.10E+08 | 52 | 0 | 1 | 9 | 26 | 6 | 1.00 | 1.00 |
| 0.2882 | 1.46E+08 | 45 | 0 | 0 | 14 | 22.5 | 6 | 0.98 | 1.00 |
| 0.2366 | 1.72E+08 | 78 | 0 | 2 | 17 | 39 | 6 | 0.97 | 1.00 |
| 0.2422 | 1.40E+08 | 61 | 0 | 2 | 16 | 30.5 | 7 | 0.85 | 1.00 |
| 0.2438 | 1.40E+08 | 53 | 0 | 1 | 14 | 26.5 | 6 | 0.98 | 1.00 |
| 0.236 | 2.17E+07 | 51 | 0 | 1 | 20 | 25.5 | 7 | 0.90 | 1.00 |
| 0.24 | 1.93E+07 | 53 | 0 | 0 | 15 | 26.5 | 7 | 0.91 | 1.00 |
| 0.2554 | 2.44E+07 | 78 | 0 | 1 | 17 | 39 | 6 | 0.90 | 1.00 |
| 0.2186 | 5.62E+06 | 68 | 1 | 1 | 23 | 33.5 | 11 | 0.66 | 0.88 |
| 0.2584 | 1.61E+06 | 70 | 0 | 1 | 24 | 35 | 11 | 0.50 | 0.97 |
| 0.2046 | 2.14E+06 | 88 | 0 | 1 | 24 | 44 | 12 | 0.51 | 0.94 |
| 0.1706 | 6.13E+07 | 76 | 0 | 2 | 23 | 38 | 16 | 0.24 | 0.99 |
| 0.1706 | 7.98E+07 | 109 | 0 | 6 | 23 | 54.5 | 12 | 0.50 | 0.97 |
| 0.1846 | 6.78E+07 | 115 | 0 | 4 | 22 | 57.5 | 12 | 0.64 | 0.97 |
| 0.166 | 5.33E+07 | 111 | 0 | 0 | 22 | 55.5 | 10 | 0.69 | 0.96 |
| 0.1428 | 6.21E+07 | 107 | 0 | 0 | 23 | 53.5 | 11 | 0.62 | 0.93 |
| 0.1768 | 5.49E+07 | 117 | 0 | 4 | 20 | 58.5 | 12 | 0.60 | 0.96 |

| | | | | | | | | | |
|---|---|---|---|---|---|---|---|---|---|
| 0.302 | 0.00E+00 | NA | NA | NA | NA | NA | NA | NA | NA |
| 0.3182 | 0.00E+00 | NA | NA | NA | NA | NA | NA | NA | NA |
| 0.3002 | 0.00E+00 | NA | NA | NA | NA | NA | NA | NA | NA |
| 0.2422 | 2.33E+07 | 33 | 0 | 3 | 14 | 16.5 | 9 | 0.79 | 0.97 |
| 0.2598 | 2.06E+07 | 27 | 0 | 1 | 16 | 13.5 | 5 | 0.89 | 1.00 |
| 0.256 | 1.71E+07 | 29 | 0 | 2 | 19 | 14.5 | 9 | 0.93 | 1.00 |
| 0.3242 | 0.00E+00 | NA | NA | NA | NA | NA | NA | NA | NA |
| 0.3424 | 0.00E+00 | NA | NA | NA | NA | NA | NA | NA | NA |
| 0.2798 | 0.00E+00 | NA | NA | NA | NA | NA | NA | NA | NA |
| 0.206 | 2.36E+07 | 39 | 0 | 4 | 19 | 19.5 | 9 | 0.95 | 1.00 |
| 0.2128 | 1.85E+07 | 47 | 0 | 4 | 21 | 23.5 | 9 | 0.74 | 0.96 |
| 0.2102 | 6.94E+06 | 43 | 0 | 3 | 19 | 21.5 | 8 | 0.79 | 0.95 |

[redacted]

| HyphaeAreaGrowth |
|---|
| 7.5138 |
| 6.942 |
| 7.4142 |
| 4.7672 |
| 6.2775 |
| 5.0146 |
| 6.675 |
| 5.1523 |
| 3.4419 |
| 5.4128 |
| 5.0884 |
| 5.3808 |
| 9.0103 |
| 7.0699 |
| 7.8595 |
| 7.6352 |
| 5.4075 |
| 5.8035 |
| 9.0972 |
| 7.6829 |
| 7.4695 |
| 5.9745 |
| 6.2077 |
| 6.8797 |
| 6.6718 |
| 7.3493 |

| |
|---|
| 8.1788 |
| 5.2843 |
| 4.51 |
| 4.0475 |
| 3.8512 |
| 2.9484 |
| 3.742 |
| 5.7907 |
| 5.065 |
| 4.9675 |
| 8.6438 |
| 8.3856 |
| 9.3524 |
| 6.1401 |
| 5.9808 |
| 5.6908 |
| 6.5075 |
| 6.6129 |
| 6.806 |
| 6.6603 |
| 6.7198 |
| 7.4578 |
| 7.3627 |
| 6.9839 |
| 8.4527 |
| 4.6473 |
| 4.853 |
| 4.301 |
| 4.4173 |

| |
|---|
| 5.5039 |
| 5.2729 |
| 5.0731 |
| 4.8113 |
| 4.6858 |
| 7.6806 |
| 6.5979 |
| 8.3038 |
| 5.705 |
| 6.637 |
| 6.2061 |
| 6.2621 |
| 7.5502 |
| 6.8523 |
| 7.1226 |
| 6.1301 |
| 6.5673 |
| 5.8244 |
| 6.8173 |
| 7.5232 |
| 3.024 |
| 3.5572 |
| 4.9459 |
| 4.9393 |
| 3.3846 |
| 5.7821 |
| 4.8832 |
| 4.8429 |
| 6.2563 |

| | |
|---|---|
| NA | |
| NA | |
| NA | |
| | 5.6245 |
| | 6.513 |
| | 6.8001 |
| NA | |
| NA | |
| NA | |
| | 6.1811 |
| | 6.2718 |
| | 5.5837 |

**Table S3:** All measurements regarding virulence

| Individual | Strain | Medium | TreatCode | Repetition | Replicate | Dead | Survival_Time | Total_time | Censorship | Mycosis |
|---|---|---|---|---|---|---|---|---|---|---|
| 1 | Control | Control | CC | 1 | 1 | 0 | 10 | 10 | 0 | - |
| 2 | Control | Control | CC | 1 | 2 | 0 | 10 | 10 | 0 | - |
| 3 | Control | Control | CC | 1 | 3 | 0 | 10 | 10 | 0 | - |
| 4 | Control | Control | CC | 1 | 4 | 0 | 10 | 10 | 0 | - |
| 5 | Control | Control | CC | 1 | 5 | 0 | 10 | 10 | 0 | - |
| 6 | Control | Control | CC | 1 | 6 | 0 | 10 | 10 | 0 | - |
| 7 | Control | Control | CC | 1 | 7 | 0 | 10 | 10 | 0 | - |
| 8 | Control | Control | CC | 1 | 8 | 0 | 10 | 10 | 0 | - |
| 9 | Control | Control | CC | 1 | 9 | 0 | 10 | 10 | 0 | - |
| 10 | Control | Control | CC | 1 | 10 | 0 | 10 | 10 | 0 | - |
| 11 | KVL1230 | MediumA | 12A | 1 | 1 | 1 | 4 | 10 | 1 | 1 |
| 12 | KVL1230 | MediumA | 12A | 1 | 2 | 1 | 5 | 10 | 1 | 1 |
| 13 | KVL1230 | MediumA | 12A | 1 | 3 | 1 | 5 | 10 | 1 | 1 |
| 14 | KVL1230 | MediumA | 12A | 1 | 4 | 1 | 5 | 10 | 1 | 1 |
| 15 | KVL1230 | MediumA | 12A | 1 | 5 | 1 | 6 | 10 | 1 | 1 |
| 16 | KVL1230 | MediumA | 12A | 1 | 6 | 1 | 9 | 10 | 1 | 1 |
| 17 | KVL1230 | MediumA | 12A | 1 | 7 | 0 | 10 | 10 | 0 | - |
| 18 | KVL1230 | MediumA | 12A | 1 | 8 | 0 | 10 | 10 | 0 | - |
| 19 | KVL1230 | MediumA | 12A | 1 | 9 | 0 | 10 | 10 | 0 | - |
| 20 | KVL1230 | MediumA | 12A | 1 | 10 | 0 | 10 | 10 | 0 | - |
| 21 | KVL1230 | MediumB | 12B | 1 | 1 | 1 | 5 | 10 | 1 | 1 |
| 22 | KVL1230 | MediumB | 12B | 1 | 2 | 1 | 5 | 10 | 1 | 1 |
| 23 | KVL1230 | MediumB | 12B | 1 | 3 | 1 | 6 | 10 | 1 | 1 |
| 24 | KVL1230 | MediumB | 12B | 1 | 4 | 1 | 6 | 10 | 1 | 1 |
| 25 | KVL1230 | MediumB | 12B | 1 | 5 | 1 | 6 | 10 | 1 | 1 |
| 26 | KVL1230 | MediumB | 12B | 1 | 6 | 1 | 7 | 10 | 1 | 1 |

| | | | | | | | | | | |
|---|---|---|---|---|---|---|---|---|---|---|
| 27 | KVL1230 | MediumB | 12B | 1 | 7 | 1 | 7 | 10 | 1 | 1 |
| 28 | KVL1230 | MediumB | 12B | 1 | 8 | 0 | 10 | 10 | 0 | - |
| 29 | KVL1230 | MediumB | 12B | 1 | 9 | 0 | 10 | 10 | 0 | - |
| 30 | KVL1230 | MediumB | 12B | 1 | 10 | 0 | 10 | 10 | 0 | - |
| 31 | KVL1230 | MediumC | 12C | 1 | 1 | 1 | 6 | 10 | 1 | 1 |
| 32 | KVL1230 | MediumC | 12C | 1 | 2 | 1 | 9 | 10 | 1 | 1 |
| 33 | KVL1230 | MediumC | 12C | 1 | 3 | 0 | 10 | 10 | 0 | - |
| 34 | KVL1230 | MediumC | 12C | 1 | 4 | 0 | 10 | 10 | 0 | - |
| 35 | KVL1230 | MediumC | 12C | 1 | 5 | 0 | 10 | 10 | 0 | - |
| 36 | KVL1230 | MediumC | 12C | 1 | 6 | 0 | 10 | 10 | 0 | - |
| 37 | KVL1230 | MediumC | 12C | 1 | 7 | 0 | 10 | 10 | 0 | - |
| 38 | KVL1230 | MediumC | 12C | 1 | 8 | 0 | 10 | 10 | 0 | - |
| 39 | KVL1230 | MediumC | 12C | 1 | 9 | 0 | 10 | 10 | 0 | - |
| 40 | KVL1230 | MediumC | 12C | 1 | 10 | 0 | 10 | 10 | 0 | 0 |
| 41 | KVL1230 | MediumD | 12D | 1 | 1 | 1 | 2 | 10 | 1 | 1 |
| 42 | KVL1230 | MediumD | 12D | 1 | 2 | 1 | 5 | 10 | 1 | 1 |
| 43 | KVL1230 | MediumD | 12D | 1 | 3 | 1 | 6 | 10 | 1 | - |
| 44 | KVL1230 | MediumD | 12D | 1 | 4 | 0 | 10 | 10 | 0 | - |
| 45 | KVL1230 | MediumD | 12D | 1 | 5 | 0 | 10 | 10 | 0 | - |
| 46 | KVL1230 | MediumD | 12D | 1 | 6 | 0 | 10 | 10 | 0 | - |
| 47 | KVL1230 | MediumD | 12D | 1 | 7 | 0 | 10 | 10 | 0 | - |
| 48 | KVL1230 | MediumD | 12D | 1 | 8 | 0 | 10 | 10 | 0 | - |
| 49 | KVL1230 | MediumD | 12D | 1 | 9 | 0 | 10 | 10 | 0 | - |
| 50 | KVL1230 | MediumD | 12D | 1 | 10 | 0 | 10 | 10 | 0 | - |
| 51 | KVL1939 | MediumA | 19A | 1 | 1 | 1 | 5 | 10 | 1 | 1 |
| 52 | KVL1939 | MediumA | 19A | 1 | 2 | 1 | 5 | 10 | 1 | 1 |
| 53 | KVL1939 | MediumA | 19A | 1 | 3 | 1 | 5 | 10 | 1 | 1 |
| 54 | KVL1939 | MediumA | 19A | 1 | 4 | 1 | 6 | 10 | 1 | 1 |
| 55 | KVL1939 | MediumA | 19A | 1 | 5 | 1 | 9 | 10 | 1 | 1 |

| | | | | | | | | | | |
|---|---|---|---|---|---|---|---|---|---|---|
| 56 | KVL1939 | MediumA | 19A | 1 | 6 | 1 | 10 | 10 | 1 | 1 |
| 57 | KVL1939 | MediumA | 19A | 1 | 7 | 0 | 10 | 10 | 0 | - |
| 58 | KVL1939 | MediumA | 19A | 1 | 8 | 0 | 10 | 10 | 0 | - |
| 59 | KVL1939 | MediumA | 19A | 1 | 9 | 0 | 10 | 10 | 0 | - |
| 60 | KVL1939 | MediumA | 19A | 1 | 10 | 0 | 10 | 10 | 0 | - |
| 61 | KVL1939 | MediumB | 19B | 1 | 1 | 1 | 5 | 10 | 1 | 1 |
| 62 | KVL1939 | MediumB | 19B | 1 | 2 | 1 | 5 | 10 | 1 | 1 |
| 63 | KVL1939 | MediumB | 19B | 1 | 3 | 1 | 6 | 10 | 1 | 1 |
| 64 | KVL1939 | MediumB | 19B | 1 | 4 | 1 | 10 | 10 | 1 | 1 |
| 65 | KVL1939 | MediumB | 19B | 1 | 5 | 0 | 10 | 10 | 0 | - |
| 66 | KVL1939 | MediumB | 19B | 1 | 6 | 0 | 10 | 10 | 0 | - |
| 67 | KVL1939 | MediumB | 19B | 1 | 7 | 0 | 10 | 10 | 0 | - |
| 68 | KVL1939 | MediumB | 19B | 1 | 8 | 0 | 10 | 10 | 0 | - |
| 69 | KVL1939 | MediumB | 19B | 1 | 9 | 0 | 10 | 10 | 0 | - |
| 70 | KVL1939 | MediumB | 19B | 1 | 10 | 0 | 10 | 10 | 0 | - |
| 71 | KVL1939 | MediumC | 19C | 1 | 1 | 1 | 5 | 10 | 1 | 0 |
| 72 | KVL1939 | MediumC | 19C | 1 | 2 | 1 | 5 | 10 | 1 | 1 |
| 73 | KVL1939 | MediumC | 19C | 1 | 3 | 1 | 5 | 10 | 1 | 1 |
| 74 | KVL1939 | MediumC | 19C | 1 | 4 | 1 | 6 | 10 | 1 | 1 |
| 75 | KVL1939 | MediumC | 19C | 1 | 5 | 1 | 6 | 10 | 1 | 0 |
| 76 | KVL1939 | MediumC | 19C | 1 | 6 | 0 | 10 | 10 | 0 | - |
| 77 | KVL1939 | MediumC | 19C | 1 | 7 | 0 | 10 | 10 | 0 | - |
| 78 | KVL1939 | MediumC | 19C | 1 | 8 | 0 | 10 | 10 | 0 | - |
| 79 | KVL1939 | MediumC | 19C | 1 | 9 | 0 | 10 | 10 | 0 | - |
| 80 | KVL1939 | MediumC | 19C | 1 | 10 | 0 | 10 | 10 | 0 | - |
| 81 | KVL1939 | MediumD | 19D | 1 | 1 | 1 | 1 | 10 | 1 | 0 |
| 82 | KVL1939 | MediumD | 19D | 1 | 2 | 1 | 4 | 10 | 1 | 1 |
| 83 | KVL1939 | MediumD | 19D | 1 | 3 | 1 | 4 | 10 | 1 | 1 |
| 84 | KVL1939 | MediumD | 19D | 1 | 4 | 1 | 6 | 10 | 1 | 1 |

| | | | | | | | | | | |
|---|---|---|---|---|---|---|---|---|---|---|
| 85 | KVL1939 | MediumD | 19D | 1 | 5 | 1 | 7 | 10 | 1 | 0 |
| 86 | KVL1939 | MediumD | 19D | 1 | 6 | 1 | 10 | 10 | 1 | 1 |
| 87 | KVL1939 | MediumD | 19D | 1 | 7 | 0 | 10 | 10 | 0 | - |
| 88 | KVL1939 | MediumD | 19D | 1 | 8 | 0 | 10 | 10 | 0 | - |
| 89 | KVL1939 | MediumD | 19D | 1 | 9 | 0 | 10 | 10 | 0 | - |
| 90 | KVL1939 | MediumD | 19D | 1 | 10 | 0 | 10 | 10 | 0 | - |
| 91 | Control | Control | CC | 2 | 1 | 0 | 10 | 10 | 0 | - |
| 92 | Control | Control | CC | 2 | 2 | 0 | 10 | 10 | 0 | - |
| 93 | Control | Control | CC | 2 | 3 | 0 | 10 | 10 | 0 | - |
| 94 | Control | Control | CC | 2 | 4 | 0 | 10 | 10 | 0 | - |
| 95 | Control | Control | CC | 2 | 5 | 0 | 10 | 10 | 0 | - |
| 96 | Control | Control | CC | 2 | 6 | 0 | 10 | 10 | 0 | - |
| 97 | Control | Control | CC | 2 | 7 | 0 | 10 | 10 | 0 | - |
| 98 | Control | Control | CC | 2 | 8 | 0 | 10 | 10 | 0 | - |
| 99 | Control | Control | CC | 2 | 9 | 0 | 10 | 10 | 0 | - |
| 100 | Control | Control | CC | 2 | 10 | 0 | 10 | 10 | 0 | - |
| 101 | KVL1230 | MediumA | 12A | 2 | 1 | 1 | 4 | 10 | 1 | 0 |
| 102 | KVL1230 | MediumA | 12A | 2 | 2 | 1 | 5 | 10 | 1 | 1 |
| 103 | KVL1230 | MediumA | 12A | 2 | 3 | 1 | 6 | 10 | 1 | 1 |
| 104 | KVL1230 | MediumA | 12A | 2 | 4 | 1 | 6 | 10 | 1 | 1 |
| 105 | KVL1230 | MediumA | 12A | 2 | 5 | 0 | 10 | 10 | 0 | - |
| 106 | KVL1230 | MediumA | 12A | 2 | 6 | 0 | 10 | 10 | 0 | - |
| 107 | KVL1230 | MediumA | 12A | 2 | 7 | 0 | 10 | 10 | 0 | - |
| 108 | KVL1230 | MediumA | 12A | 2 | 8 | 0 | 10 | 10 | 0 | - |
| 109 | KVL1230 | MediumA | 12A | 2 | 9 | 0 | 10 | 10 | 0 | - |
| 110 | KVL1230 | MediumA | 12A | 2 | 10 | 0 | 10 | 10 | 0 | - |
| 111 | KVL1230 | MediumB | 12B | 2 | 1 | 1 | 2 | 10 | 1 | 0 |
| 112 | KVL1230 | MediumB | 12B | 2 | 2 | 1 | 6 | 10 | 1 | 1 |
| 113 | KVL1230 | MediumB | 12B | 2 | 3 | 1 | 6 | 10 | 1 | 1 |

| | | | | | | | | | | |
|---|---|---|---|---|---|---|---|---|---|---|
| 114 | KVL1230 | MediumB | 12B | 2 | 4 | 1 | 6 | 10 | 1 | 1 |
| 115 | KVL1230 | MediumB | 12B | 2 | 5 | 1 | 6 | 10 | 1 | 1 |
| 116 | KVL1230 | MediumB | 12B | 2 | 6 | 1 | 8 | 10 | 1 | 1 |
| 117 | KVL1230 | MediumB | 12B | 2 | 7 | 1 | 9 | 10 | 1 | 1 |
| 118 | KVL1230 | MediumB | 12B | 2 | 8 | 1 | 9 | 10 | 1 | 1 |
| 119 | KVL1230 | MediumB | 12B | 2 | 9 | 0 | 10 | 10 | 0 | - |
| 120 | KVL1230 | MediumB | 12B | 2 | 10 | 0 | 10 | 10 | 0 | - |
| 121 | KVL1230 | MediumC | 12C | 2 | 1 | 1 | 6 | 10 | 1 | 1 |
| 122 | KVL1230 | MediumC | 12C | 2 | 2 | 1 | 6 | 10 | 1 | 1 |
| 123 | KVL1230 | MediumC | 12C | 2 | 3 | 0 | 10 | 10 | 0 | - |
| 124 | KVL1230 | MediumC | 12C | 2 | 4 | 0 | 10 | 10 | 0 | - |
| 125 | KVL1230 | MediumC | 12C | 2 | 5 | 0 | 10 | 10 | 0 | - |
| 126 | KVL1230 | MediumC | 12C | 2 | 6 | 0 | 10 | 10 | 0 | - |
| 127 | KVL1230 | MediumC | 12C | 2 | 7 | 0 | 10 | 10 | 0 | - |
| 128 | KVL1230 | MediumC | 12C | 2 | 8 | 0 | 10 | 10 | 0 | - |
| 129 | KVL1230 | MediumC | 12C | 2 | 9 | 0 | 10 | 10 | 0 | - |
| 130 | KVL1230 | MediumC | 12C | 2 | 10 | 0 | 10 | 10 | 0 | - |
| 131 | KVL1230 | MediumD | 12D | 2 | 1 | 1 | 6 | 10 | 1 | 1 |
| 132 | KVL1230 | MediumD | 12D | 2 | 2 | 1 | 6 | 10 | 1 | 1 |
| 133 | KVL1230 | MediumD | 12D | 2 | 3 | 1 | 8 | 10 | 1 | 1 |
| 134 | KVL1230 | MediumD | 12D | 2 | 4 | 0 | 10 | 10 | 0 | - |
| 135 | KVL1230 | MediumD | 12D | 2 | 5 | 0 | 10 | 10 | 0 | - |
| 136 | KVL1230 | MediumD | 12D | 2 | 6 | 0 | 10 | 10 | 0 | - |
| 137 | KVL1230 | MediumD | 12D | 2 | 7 | 0 | 10 | 10 | 0 | - |
| 138 | KVL1230 | MediumD | 12D | 2 | 8 | 0 | 10 | 10 | 0 | - |
| 139 | KVL1230 | MediumD | 12D | 2 | 9 | 0 | 10 | 10 | 0 | - |
| 140 | KVL1230 | MediumD | 12D | 2 | 10 | 0 | 10 | 10 | 0 | - |
| 141 | KVL1939 | MediumA | 19A | 2 | 1 | 1 | 1 | 10 | 1 | 0 |
| 142 | KVL1939 | MediumA | 19A | 2 | 2 | 1 | 4 | 10 | 1 | 1 |

| | | | | | | | | | | |
|---|---|---|---|---|---|---|---|---|---|---|
| 143 | KVL1939 | MediumA | 19A | 2 | 3 | 1 | 6 | 10 | 1 | 1 |
| 144 | KVL1939 | MediumA | 19A | 2 | 4 | 1 | 6 | 10 | 1 | 1 |
| 145 | KVL1939 | MediumA | 19A | 2 | 5 | 1 | 9 | 10 | 1 | 0 |
| 146 | KVL1939 | MediumA | 19A | 2 | 6 | 1 | 10 | 10 | 1 | 1 |
| 147 | KVL1939 | MediumA | 19A | 2 | 7 | 0 | 10 | 10 | 0 | - |
| 148 | KVL1939 | MediumA | 19A | 2 | 8 | 0 | 10 | 10 | 0 | - |
| 149 | KVL1939 | MediumA | 19A | 2 | 9 | 0 | 10 | 10 | 0 | - |
| 150 | KVL1939 | MediumA | 19A | 2 | 10 | 0 | 10 | 10 | 0 | - |
| 151 | KVL1939 | MediumB | 19B | 2 | 1 | 1 | 5 | 10 | 1 | 1 |
| 152 | KVL1939 | MediumB | 19B | 2 | 2 | 1 | 5 | 10 | 1 | 1 |
| 153 | KVL1939 | MediumB | 19B | 2 | 3 | 1 | 6 | 10 | 1 | 1 |
| 154 | KVL1939 | MediumB | 19B | 2 | 4 | 1 | 10 | 10 | 1 | 1 |
| 155 | KVL1939 | MediumB | 19B | 2 | 5 | 0 | 10 | 10 | 0 | - |
| 156 | KVL1939 | MediumB | 19B | 2 | 6 | 0 | 10 | 10 | 0 | - |
| 157 | KVL1939 | MediumB | 19B | 2 | 7 | 0 | 10 | 10 | 0 | - |
| 158 | KVL1939 | MediumB | 19B | 2 | 8 | 0 | 10 | 10 | 0 | - |
| 159 | KVL1939 | MediumB | 19B | 2 | 9 | 0 | 10 | 10 | 0 | - |
| 160 | KVL1939 | MediumB | 19B | 2 | 10 | 0 | 10 | 10 | 0 | - |
| 161 | KVL1939 | MediumC | 19C | 2 | 1 | 1 | 5 | 10 | 1 | 1 |
| 162 | KVL1939 | MediumC | 19C | 2 | 2 | 1 | 8 | 10 | 1 | 1 |
| 163 | KVL1939 | MediumC | 19C | 2 | 3 | 1 | 10 | 10 | 1 | 1 |
| 164 | KVL1939 | MediumC | 19C | 2 | 4 | 0 | 10 | 10 | 0 | - |
| 165 | KVL1939 | MediumC | 19C | 2 | 5 | 0 | 10 | 10 | 0 | - |
| 166 | KVL1939 | MediumC | 19C | 2 | 6 | 0 | 10 | 10 | 0 | - |
| 167 | KVL1939 | MediumC | 19C | 2 | 7 | 0 | 10 | 10 | 0 | - |
| 168 | KVL1939 | MediumC | 19C | 2 | 8 | 0 | 10 | 10 | 0 | - |
| 169 | KVL1939 | MediumC | 19C | 2 | 9 | 0 | 10 | 10 | 0 | - |
| 170 | KVL1939 | MediumC | 19C | 2 | 10 | 0 | 10 | 10 | 0 | - |
| 171 | KVL1939 | MediumD | 19D | 2 | 1 | 0 | 10 | 10 | 0 | - |

| | | | | | | | | | | |
|---|---|---|---|---|---|---|---|---|---|---|
| 172 | KVL1939 | MediumD | 19D | 2 | 2 | 0 | 10 | 10 | 0 | - |
| 173 | KVL1939 | MediumD | 19D | 2 | 3 | 0 | 10 | 10 | 0 | - |
| 174 | KVL1939 | MediumD | 19D | 2 | 4 | 0 | 10 | 10 | 0 | - |
| 175 | KVL1939 | MediumD | 19D | 2 | 5 | 0 | 10 | 10 | 0 | - |
| 176 | KVL1939 | MediumD | 19D | 2 | 6 | 0 | 10 | 10 | 0 | - |
| 177 | KVL1939 | MediumD | 19D | 2 | 7 | 0 | 10 | 10 | 0 | - |
| 178 | KVL1939 | MediumD | 19D | 2 | 8 | 0 | 10 | 10 | 0 | - |
| 179 | KVL1939 | MediumD | 19D | 2 | 9 | 0 | 10 | 10 | 0 | - |
| 180 | KVL1939 | MediumD | 19D | 2 | 10 | 0 | 10 | 10 | 0 | - |
| 181 | Control | Control | CC | 3 | 1 | 0 | 10 | 10 | 0 | - |
| 182 | Control | Control | CC | 3 | 2 | 0 | 10 | 10 | 0 | - |
| 183 | Control | Control | CC | 3 | 3 | 0 | 10 | 10 | 0 | - |
| 184 | Control | Control | CC | 3 | 4 | 0 | 10 | 10 | 0 | - |
| 185 | Control | Control | CC | 3 | 5 | 0 | 10 | 10 | 0 | - |
| 186 | Control | Control | CC | 3 | 6 | 0 | 10 | 10 | 0 | - |
| 187 | Control | Control | CC | 3 | 7 | 0 | 10 | 10 | 0 | - |
| 188 | Control | Control | CC | 3 | 8 | 0 | 10 | 10 | 0 | - |
| 189 | Control | Control | CC | 3 | 9 | 0 | 10 | 10 | 0 | - |
| 190 | Control | Control | CC | 3 | 10 | 0 | 10 | 10 | 0 | - |
| 191 | KVL1230 | MediumA | 12A | 3 | 1 | 1 | 1 | 10 | 1 | 0 |
| 192 | KVL1230 | MediumA | 12A | 3 | 2 | 1 | 5 | 10 | 1 | 1 |
| 193 | KVL1230 | MediumA | 12A | 3 | 3 | 1 | 5 | 10 | 1 | 1 |
| 194 | KVL1230 | MediumA | 12A | 3 | 4 | 1 | 5 | 10 | 1 | 1 |
| 195 | KVL1230 | MediumA | 12A | 3 | 5 | 1 | 8 | 10 | 1 | 1 |
| 196 | KVL1230 | MediumA | 12A | 3 | 6 | 0 | 10 | 10 | 0 | - |
| 197 | KVL1230 | MediumA | 12A | 3 | 7 | 0 | 10 | 10 | 0 | - |
| 198 | KVL1230 | MediumA | 12A | 3 | 8 | 0 | 10 | 10 | 0 | - |
| 199 | KVL1230 | MediumA | 12A | 3 | 9 | 0 | 10 | 10 | 0 | - |
| 200 | KVL1230 | MediumA | 12A | 3 | 10 | 0 | 10 | 10 | 0 | - |

| | | | | | | | | | | |
|---|---|---|---|---|---|---|---|---|---|---|
| 201 | KVL1230 | MediumB | 12B | 3 | 1 | 1 | 5 | 10 | 1 | 1 |
| 202 | KVL1230 | MediumB | 12B | 3 | 2 | 1 | 5 | 10 | 1 | 1 |
| 203 | KVL1230 | MediumB | 12B | 3 | 3 | 1 | 5 | 10 | 1 | 1 |
| 204 | KVL1230 | MediumB | 12B | 3 | 4 | 1 | 7 | 10 | 1 | 1 |
| 205 | KVL1230 | MediumB | 12B | 3 | 5 | 1 | 9 | 10 | 1 | 1 |
| 206 | KVL1230 | MediumB | 12B | 3 | 6 | 0 | 10 | 10 | 0 | - |
| 207 | KVL1230 | MediumB | 12B | 3 | 7 | 0 | 10 | 10 | 0 | - |
| 208 | KVL1230 | MediumB | 12B | 3 | 8 | 0 | 10 | 10 | 0 | - |
| 209 | KVL1230 | MediumB | 12B | 3 | 9 | 0 | 10 | 10 | 0 | - |
| 210 | KVL1230 | MediumB | 12B | 3 | 10 | 0 | 10 | 10 | 0 | - |
| 211 | KVL1230 | MediumC | 12C | 3 | 1 | 1 | 1 | 10 | 1 | 0 |
| 212 | KVL1230 | MediumC | 12C | 3 | 2 | 1 | 6 | 10 | 1 | 1 |
| 213 | KVL1230 | MediumC | 12C | 3 | 3 | 1 | 8 | 10 | 1 | 1 |
| 214 | KVL1230 | MediumC | 12C | 3 | 4 | 1 | 10 | 10 | 1 | 1 |
| 215 | KVL1230 | MediumC | 12C | 3 | 5 | 0 | 10 | 10 | 0 | - |
| 216 | KVL1230 | MediumC | 12C | 3 | 6 | 0 | 10 | 10 | 0 | - |
| 217 | KVL1230 | MediumC | 12C | 3 | 7 | 0 | 10 | 10 | 0 | - |
| 218 | KVL1230 | MediumC | 12C | 3 | 8 | 0 | 10 | 10 | 0 | - |
| 219 | KVL1230 | MediumC | 12C | 3 | 9 | 0 | 10 | 10 | 0 | - |
| 220 | KVL1230 | MediumC | 12C | 3 | 10 | 0 | 10 | 10 | 0 | - |
| 221 | KVL1230 | MediumD | 12D | 3 | 1 | 1 | 8 | 10 | 1 | 1 |
| 222 | KVL1230 | MediumD | 12D | 3 | 2 | 0 | 10 | 10 | 0 | - |
| 223 | KVL1230 | MediumD | 12D | 3 | 3 | 0 | 10 | 10 | 0 | - |
| 224 | KVL1230 | MediumD | 12D | 3 | 4 | 0 | 10 | 10 | 0 | - |
| 225 | KVL1230 | MediumD | 12D | 3 | 5 | 0 | 10 | 10 | 0 | - |
| 226 | KVL1230 | MediumD | 12D | 3 | 6 | 0 | 10 | 10 | 0 | - |
| 227 | KVL1230 | MediumD | 12D | 3 | 7 | 0 | 10 | 10 | 0 | - |
| 228 | KVL1230 | MediumD | 12D | 3 | 8 | 0 | 10 | 10 | 0 | - |
| 229 | KVL1230 | MediumD | 12D | 3 | 9 | 0 | 10 | 10 | 0 | - |

| | | | | | | | | | | |
|---|---|---|---|---|---|---|---|---|---|---|
| 230 | KVL1230 | MediumD | 12D | 3 | 10 | 0 | 10 | 10 | 0 | - |
| 231 | KVL1939 | MediumA | 19A | 3 | 1 | 1 | 1 | 10 | 1 | 0 |
| 232 | KVL1939 | MediumA | 19A | 3 | 2 | 1 | 2 | 10 | 1 | 0 |
| 233 | KVL1939 | MediumA | 19A | 3 | 3 | 1 | 6 | 10 | 1 | 1 |
| 234 | KVL1939 | MediumA | 19A | 3 | 4 | 0 | 10 | 10 | 0 | - |
| 235 | KVL1939 | MediumA | 19A | 3 | 5 | 0 | 10 | 10 | 0 | - |
| 236 | KVL1939 | MediumA | 19A | 3 | 6 | 0 | 10 | 10 | 0 | - |
| 237 | KVL1939 | MediumA | 19A | 3 | 7 | 0 | 10 | 10 | 0 | - |
| 238 | KVL1939 | MediumA | 19A | 3 | 8 | 0 | 10 | 10 | 0 | - |
| 239 | KVL1939 | MediumA | 19A | 3 | 9 | 0 | 10 | 10 | 0 | - |
| 240 | KVL1939 | MediumA | 19A | 3 | 10 | 0 | 10 | 10 | 0 | - |
| 241 | KVL1939 | MediumB | 19B | 3 | 1 | 1 | 1 | 10 | 1 | 0 |
| 242 | KVL1939 | MediumB | 19B | 3 | 2 | 1 | 2 | 10 | 1 | 1 |
| 243 | KVL1939 | MediumB | 19B | 3 | 3 | 1 | 4 | 10 | 1 | 0 |
| 244 | KVL1939 | MediumB | 19B | 3 | 4 | 1 | 8 | 10 | 1 | 1 |
| 245 | KVL1939 | MediumB | 19B | 3 | 5 | 1 | 8 | 10 | 1 | 1 |
| 246 | KVL1939 | MediumB | 19B | 3 | 6 | 0 | 10 | 10 | 0 | - |
| 247 | KVL1939 | MediumB | 19B | 3 | 7 | 0 | 10 | 10 | 0 | - |
| 248 | KVL1939 | MediumB | 19B | 3 | 8 | 0 | 10 | 10 | 0 | - |
| 249 | KVL1939 | MediumB | 19B | 3 | 9 | 0 | 10 | 10 | 0 | - |
| 250 | KVL1939 | MediumB | 19B | 3 | 10 | 0 | 10 | 10 | 0 | - |
| 251 | KVL1939 | MediumC | 19C | 3 | 1 | 1 | 5 | 10 | 1 | 1 |
| 252 | KVL1939 | MediumC | 19C | 3 | 2 | 1 | 7 | 10 | 1 | 1 |
| 253 | KVL1939 | MediumC | 19C | 3 | 3 | 0 | 10 | 10 | 0 | - |
| 254 | KVL1939 | MediumC | 19C | 3 | 4 | 0 | 10 | 10 | 0 | - |
| 255 | KVL1939 | MediumC | 19C | 3 | 5 | 0 | 10 | 10 | 0 | - |
| 256 | KVL1939 | MediumC | 19C | 3 | 6 | 0 | 10 | 10 | 0 | - |
| 257 | KVL1939 | MediumC | 19C | 3 | 7 | 0 | 10 | 10 | 0 | - |
| 258 | KVL1939 | MediumC | 19C | 3 | 8 | 0 | 10 | 10 | 0 | - |

| | | | | | | | | | | |
|---|---|---|---|---|---|---|---|---|---|---|
| 259 | KVL1939 | MediumC | 19C | 3 | 9 | 0 | 10 | 10 | 0 | - |
| 260 | KVL1939 | MediumC | 19C | 3 | 10 | 0 | 10 | 10 | 0 | - |
| 261 | KVL1939 | MediumD | 19D | 3 | 1 | 1 | 5 | 10 | 1 | 1 |
| 262 | KVL1939 | MediumD | 19D | 3 | 2 | 1 | 6 | 10 | 1 | 1 |
| 263 | KVL1939 | MediumD | 19D | 3 | 3 | 1 | 6 | 10 | 1 | 0 |
| 264 | KVL1939 | MediumD | 19D | 3 | 4 | 1 | 8 | 10 | 1 | 1 |
| 265 | KVL1939 | MediumD | 19D | 3 | 5 | 1 | 10 | 10 | 1 | 1 |
| 266 | KVL1939 | MediumD | 19D | 3 | 6 | 0 | 10 | 10 | 0 | - |
| 267 | KVL1939 | MediumD | 19D | 3 | 7 | 0 | 10 | 10 | 0 | - |
| 268 | KVL1939 | MediumD | 19D | 3 | 8 | 0 | 10 | 10 | 0 | - |
| 269 | KVL1939 | MediumD | 19D | 3 | 9 | 0 | 10 | 10 | 0 | - |
| 270 | KVL1939 | MediumD | 19D | 3 | 10 | 0 | 10 | 10 | 0 | - |

| Comments |
| --- |
| Photo |
| Photo |

| |
|---|
| Photo |
| Photo |
| Photo |

| |
|---|
| Photo |
| Photo |
| Photo |
